# Representational structure or task structure? Bias in neural representational similarity analysis and a Bayesian method for reducing bias

**DOI:** 10.1101/347260

**Authors:** Ming Bo Cai, Nicolas W. Schuck, Jonathan W. Pillow, Yael Niv

## Abstract

The activity of neural populations in the brains of humans and animals can exhibit vastly different spatial patterns when faced with different tasks or environmental stimuli. The degree of similarity between these neural activity patterns in response to different events is used to characterize the representational structure of cognitive states in a neural population. The dominant methods of investigating this similarity structure first estimate neural activity patterns from noisy neural imaging data using linear regression, and then examine the similarity between the estimated patterns. Here, we show that this approach introduces spurious bias structure in the resulting similarity matrix, in particular when applied to fMRI data. This problem is especially severe when the signal-to-noise ratio is low and in cases where experimental conditions cannot be fully randomized in a task. We propose Bayesian Representational Similarity Analysis (BRSA), an alternative method for computing representational similarity, in which we treat the covariance structure of neural activity patterns as a hyper-parameter in a generative model of the neural data. By marginalizing over the unknown activity patterns, we can directly estimate this covariance structure from imaging data. This method offers significant reductions in bias and allows estimation of neural representational similarity with previously unattained levels of precision at low signal-to-noise ratio. The probabilistic framework allows for jointly analyzing data from a group of participants. The method can also simultaneously estimate a signal-to-noise ratio map that shows where the learned representational structure is supported more strongly. Both this map and the learned covariance matrix can be used as a structured prior for maximum *a posteriori* estimation of neural activity patterns, which can be further used for fMRI decoding. We make our tool freely available in Brain Imaging Analysis Kit (BrainIAK).

**Author summary:** We show the severity of the bias introduced when performing representational similarity analysis (RSA) based on neural activity pattern estimated within imaging runs. Our Bayesian RSA method significantly reduces the bias and can learn a shared representational structure across multiple participants. We also demonstrate its extension as a new multi-class decoding tool.

## Introduction

Functional magnetic resonance imaging (fMRI) measures the blood-oxygen-level-dependent (BOLD) signals [1], which rise to peak ∼ 6 seconds after neuronal activity increases in a local region [2]. Because of its non-invasiveness, full-brain coverage, and relatively favorable trade-off between spatial and temporal resolution, fMRI has been a powerful tool to study the neural correlates of cognition [3-5]. In the last decade, research has moved beyond simply localizing the brain regions selectively activated by the cognitive processes and focus has been increasingly placed on the relationship between the detailed spatial patterns of neural activity and cognitive processes [6, 7].

An important tool for characterizing the functional architecture of sensory cortex is representational similarity analysis (RSA) [8]. This classic method first estimates the neural activity pattern from fMRI data recorded as participants observe a set of stimuli or experience a set of task conditions, and then calculates the similarity (e.g., by Pearson correlation) between each pair of the estimated patterns. The rationale is that if two stimuli are represented with similar codes in a brain region, the spatial patterns of neural activation in that region would be similar when processing these two stimuli.

After the similarity matrix between all pairs of activity patterns is calculated in an ROI, it can be compared against similarity matrices predicted by candidate computational models. Researchers can also convert the similarity matrix into a representational dissimilarity matrix (RDM, e.g., 1 − *C*, for similarity *C* based on correlation) and visualize the structure of the representational space in the ROI by projecting the dissimilarity matrix to a low dimensional space [8]. Researchers might also test whether certain experimental manipulations changes the degrees of similarity between neural patterns of interest [9, 10]. To list just a few application of this method in the domain of visual neuroscience, RSA has revealed that humans and monkeys have highly similar representational structures in the inferotemporal (IT) cortex for images across various semantic categories [11]. It also revealed a continuum in the abstract representation of biological classes in human ventral object visual cortex [12] and that basic categorical structure gradually emerges through the hierarchy of visual cortex [13]. Because of the additional flexibility of exploring the structure of neural representation without building explicit computational models, RSA has also gained popularity among cognitive neuroscientists for studying more complex tasks beyond perception, such as decision making.

While RSA has been widely adopted in many fields of cognitive neuroscience, a few recent studies have revealed that the similarity structure estimated by standard RSA might be confounded by various factors. First, the calculated similarity between two neural patterns strongly depends on the time that elapsed between the two measured patterns: the closer the two patterns are in time, the more similar they are [14] [15]. Second, it was found that because different brain regions share some common time course of fluctuation independent of the stimuli being presented (intrinsic fluctuations), RDMs between regions are highly similar when calculated based on patterns of the same trials of tasks but not when they are calculated based on separate trials (thus the intrinsic fluctuation are not shared across regions). This indicates that RSA can be strongly influenced by intrinsic fluctuation [14]. Lastly, Diedrichsen et al. (2011) pointed out that the noise in the estimated activity patterns can add a diagonal component to the condition-by-condition covariance matrix of the spatial patterns. This leads to over-estimation of the variance of the neural pattern and underestimation of correlation between true patterns, and this underestimation depends on signal-to-noise ratio in each ROI, making it difficult to make comparison of RDMs between regions [16].

Recognizing the first two issues, several groups have recently suggested modifications to RSA such as calculating similarity or distance between activity patterns estimated from separate fMRI runs [15, 17], henceforth referred to as cross-run RSA, and using a Taylor expansion to approximate and regress out the dependency of pattern similarity on the interval between events [15]. For the last issue, Diedrichsen et al. (2011) proposed modeling the condition-by-condition covariance matrix between estimated neural patterns as the sum of a diagonal component that models the contribution of noise in the estimated neural patterns to the covariance matrix and components reflecting the researcher’s hypothetical representational structure in the ROI [16] (“pattern-component model”; PCM). These methods improve on traditional RSA, but are not explicitly directed at the source of the bias, and therefore only offer partial solutions.

Indeed, the severity of confounds in traditional RSA is not yet widely recognized. RSA based on neural patterns estimated within an imaging run is still commonly performed. Furthermore, sometimes a study might need to examine the representational similarity between task conditions within an imaging run, such that cross-run RSA is not feasible. The Taylor expansion approach to model the effect of event-interval can be difficult to set up when a task condition repeats several times in an experiment. There also lacks a detailed mathematical examination of the source of the bias and how different ways of applying RSA affect the bias. Researchers sometimes hold the view that RSA of raw fMRI patterns instead of activity patterns (*β*) estimated through a general linear model (GLM) [18] does not suffer from the confounds mentioned above. Last but not least, the contribution of noise in the estimated neural patterns to the sample covariance matrix between patterns may not be restricted to the diagonal elements, as we will demonstrate below.

In this paper, we first compare the result of performing traditional RSA on a task-based fMRI dataset with the results obtained when performing the same analysis on white noise, to illustrate the severe bias and spurious similarity structure that can result from that performing RSA on pattern estimates within imaging runs. By applying task-specific RSA on irrelevant resting-state fMRI data, we show that spurious structure also emerges when RSA is performed on the raw fMRI pattern rather than estimated task activation patterns. We sow that the spurious structure can be far from a diagonal matrix, and masks any true similarity structure. We then provide an analytic derivation to help understand the source of the bias in traditional RSA. Previously, we have proposed a method named Bayesian RSA (BRSA), which significantly reduced this bias and allows analysis within imaging runs [19]. Here, we further extend the method to explicitly model spatial noise correlation, thereby mitigating the second issue identified by Heriksson et al. [14], namely the intrinsic fluctuation not modelled by task events in an experiment. Furthermore, inspired by the methods of hyper-alignment [20] and shared response models [21], we extend our method to learn a shared representational similarity structure across multiple participants (Group BRSA) and demonstrate improved accuracy of this approach. Since our method significantly reduces bias in the estimated similarity matrix but does not fully eliminate it at regimes of very low signal-to-noise ratio (SNR), we further provide a cross-validation approach to detecting over-fitting to the data. Finally, we show that the learned representational structure can serve as an empirical prior to constrain the posterior estimation of activity patterns, which can be used to decode the cognitive state underlying activity observed in new fMRI data.

The algorithm in this paper is publicly available in the python package Brain Imaging Analysis Kit (BrainIAK).^1^

## Results

### Traditional RSA translates structured noise in estimated activity patterns into spurious similarity structure

Traditional RSA [8] first estimates the response amplitudes (***β***) of each voxel in an ROI and then calculates the similarity between the estimated spatial response patterns of that ROI to different task conditions.

The estimation of ***β*** is based on a GLM. We denote the fMRI time series from an experiment as 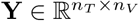, with *n_T_* being the number of time points and *n_V_* the number of voxels. The GLM assumes that
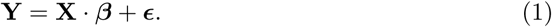

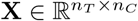 is the “design matrix,” where *n_C_* is the number of task conditions. Each column of the design matrix is constructed by convolving a hemodynamic response function (HRF) with a time series describing the onsets and duration of all events belonging to one task condition. The regressors composing the design matrix express the hypothesized response time course elicited by each task condition. Each voxel’s response amplitudes to different task conditions can differ. All voxels’ response profiles form a matrix of spatial activity patterns 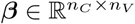, with each row representing the spatial pattern of activity elicited by one task condition. The responses to all conditions are assumed to contribute linearly to the spatio-temporal fMRI signal through the temporal profile of hemodynamic response expressed in **X**. Thus, the measured **Y** is assumed to be a linear sum of **X** weighted by response amplitude ***β***, corrupted by zero-mean noise *ϵ*.

The goal of RSA is to understand the degree of similarity between each pair of spatial response patterns (i.e., between the rows of ***β***). But because the true ***β*** is not accessible, a point estimate of ***β***, derived through linear regression, is usually used as a surrogate:
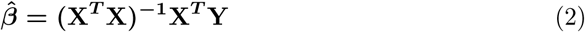

Similarity is then calculated between rows of 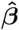. For instance, one measure of similarity that is frequently used is Pearson correlation:
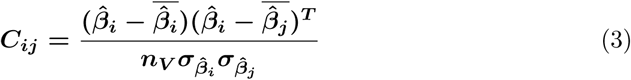

To demonstrate the spurious structure that may appear in the result of traditional RSA, we first performed RSA on the fMRI data in one ROI, the orbitofrontal cortex, in a previous dataset involving a decision-making task [22]. The task included 16 different task conditions, or “states.” In each state, participants paid attention to one of two overlapping images (face or house) and made judgments about the image in the attended category. The transition between the 16 task states followed the Markov chain shown in Fig **1A**, thus some states often preceded certain other states. The 16 states could be grouped into 3 categories according to the structure of transitions among states (the exact meaning of the states, or the 3 categories, are not important in the context of the discussion here.) We performed traditional RSA on the 16 estimated spatial response patterns corresponding to the 16 task states. To visualize the structure of the neural representation of the task states in the ROI, we used multi-dimensional scaling (MDS) [23] to project the 16-dimensional space defined by the distance between states (1 - correlation) onto a 3-dimensional space (Fig **1B**).

**Figure 1.**
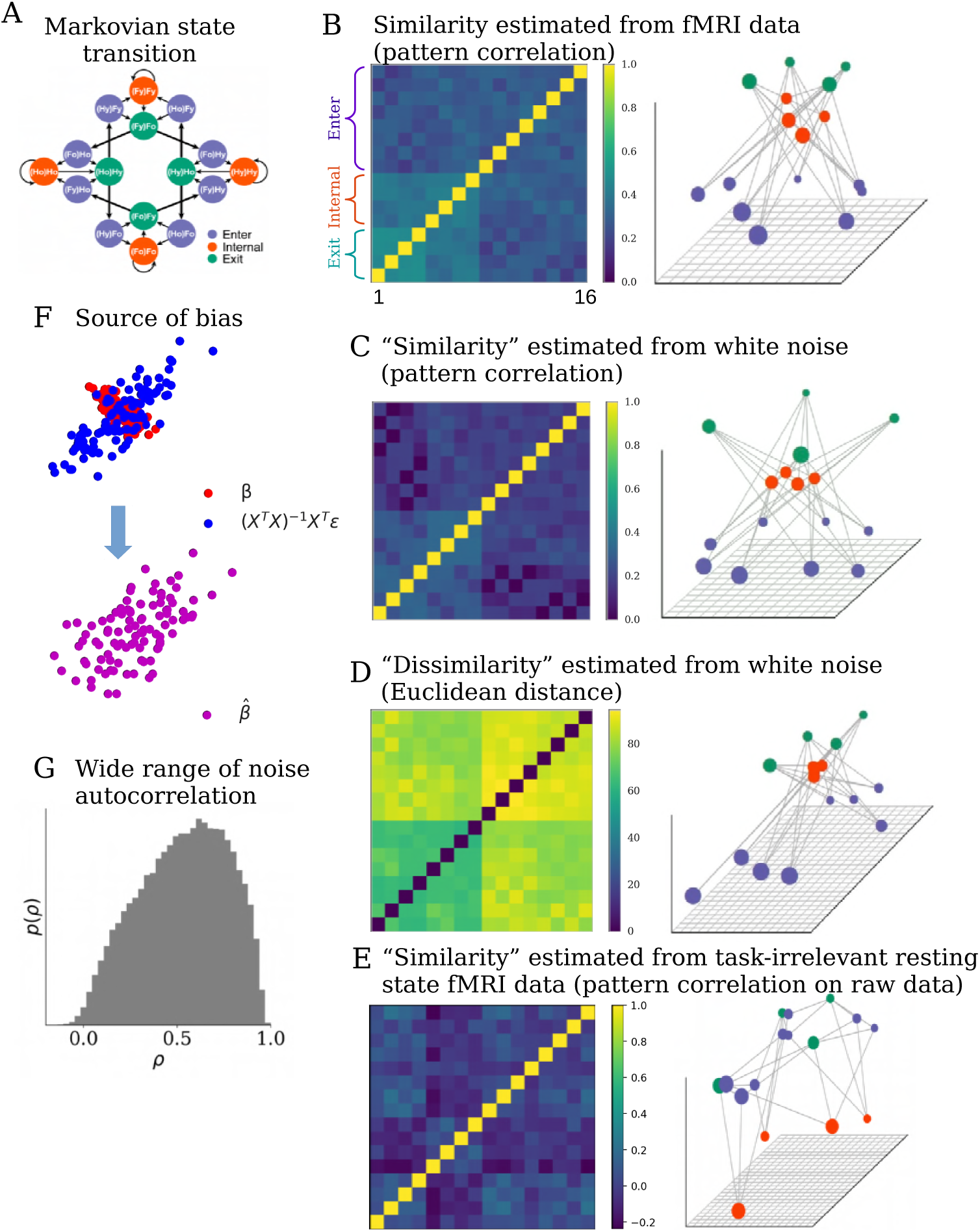
Standard RSA introduces bias structure to the similarity matrix. **(A)** A cognitive task including 16 different experimental conditions. Transitions between conditions follow a Markov process. Arrows indicate possible transitions, each with ***p* = 0.5**. The task conditions can be grouped into 3 categories (color coded) according to their characteristic transition structure. **(B)** Standard RSA of activity patterns corresponding to each condition estimated from a brain region reveals a highly structured similarity matrix (left) that reflects aspects of the transition structure in the task. Converting the similarity matrix ***C*** to a distance matrix **1 − *C*** and projecting it to a low-dimensional space using MDS reveals a highly regular structure (right). Seeing such a result, one may infer that representational structure in the ROI strongly reflects the task structure. **(C)** However, applying RSA to regression estimates of of patterns obtained from pure white noise generates a very similar similarity matrix (left), with a similar low-dimensional projection (right). This indicates that standard RSA can introduce spurious structure in the similarity matrix that does not exist in the data. **(D)** RSA Using Euclidean distance as a similarity metric applied to patterns estimated from the same noise (left) yields a slightly different, but still structured, similarity structure (right). **(E)** Calculating the correlation between raw patterns of resting state fMRI data (instead of patterns estimated by a GLM), assuming the same task structure as in (A), also generates spurious similarity structure, albeit different from those in (B-D). A permutation test shows that many of the high correlation values are not expected in a null distribution (details in main text). **(F)** The bias in this case comes from structured noise introduced during the GLM analysis. Assuming the true patterns ***β*** (red dots) of two task conditions are anti-correlated (the horizontal and vertical coordinates of each dot represent the response amplitudes of one voxel to the two task conditions), regression turns the noise ***ϵ*** in fMRI data into structured noise **(X*^T^* X)^−1^X*^T^ ϵ*** (blue dots). The correlation between the noises in the estimated patterns is often non-zero (assumed to be positive correlation here) due to the correlation structure in the design matrix and the autocorrelation property of the noise. The estimated patterns ***β*** (purple dots) are the sum of ***β*** and **(X*^T^* X)^−1^X*^T^ ϵ***. The correlation structure between estimated activity vectors for each condition will therefore differ from the correlation structure between the true patterns ***β***. **(G)** Distribution of the autocorrelation coefficients in a resting state fMRI dataset, estimated by fitting AR(1) model to the time series of each voxel resampled at TR=2.4s. The wide range of degree of autocorrelation across voxels makes it difficulty to calculate a simple analytic form of the bias structure introduced by the structured noise, and calls for modeling the noise structure of each voxel separately.

This projection appears to show clear grouping of the states in the orbitofrontal cortex consistent with the 3 categories, suggesting that this brain area represents this aspect of the task. However, a similar representational structure was also observed in other ROIs. In addition, when we applied the same GLM to randomly generated white noise and performed RSA on the resulting parameter estimates, the similarity matrix closely resembled the result found in the real fMRI data (Fig **1C**). Since there is no task-related activity in the white noise, the structure obtained from white noise is clearly spurious and must reflect a bias introduced by the analysis. In fact, we found that the off-diagonal structure obtained from white noise (Fig **1C**) explained **84 ± 12%** of the variance of the off-diagonals obtained from real data (Fig **1B**). This shows that the bias introduced by traditional RSA can dominate the result, masking the real representational structure in the data.

To help understand this observation, we provide an analytic derivation of the bias with a few simplifying assumptions [19]. The calculation of the sample correlation of 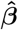 in traditional RSA implies the implicit assumption that an underlying covariance structure exists that describe the distribution of ***β***, and the activity profile of each voxel is one sample from this distribution. Therefore, examining the relation between the covariance of 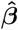 and that of true ***β*** will help us understand the bias in traditional RSA.

We assume that a covariance matrix **U** (of size ***n_C_* × *n_C_***) captures the true covariance structure of ***β*** across all voxels in the ROI: ***β*** ∼ **N(0, U)**. Similarity measures such as correlation are derived from **U** by normalizing the diagonal elements to 1. It is well known that temporal autocorrelation exists in fMRI noise [24, 25]. To capture this, we assume that in each voxel ***ϵ*** ∼ ***N* (0, Σ*_ϵ_*)**, where 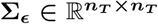 is the temporal covariance of the noise (for illustration purposes, here we assume that all voxels have the same noise variance and autocorrelation, and temporarily assume the noise is spatially independent).

By substituting the expression for **Y** from equation 1 we obtain
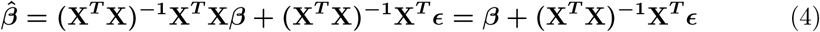

which means the point estimate of ***β*** is contaminated by a noise term **(X*^T^* X)^−1^X*^T^ ϵ***. Assuming that the signal ***β*** is independent from the noise ***ϵ***, it is then also independent from the linear transformation of the noise, **(X*^T^* X)^−1^X*^T^ ϵ***. Thus the covariance of 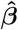 is the sum of the covariance of true ***β*** and the covariance of **(X*^T^* X)^−1^X*^T^ ϵ***:
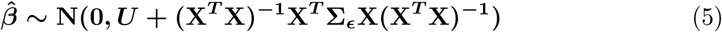

The term **(X*^T^* X)^−1^X*^T^* Σ*_ϵ_*X(X*^T^* X)^−1^** is the source of the bias in RSA. This bias originates from the structured noise **(X*^T^* X)^−1^X*^T^ ϵ*** in estimating 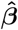. It depends on both the design matrix **X** and the temporal autocorrelation of the noise ***ϵ***. Fig **1F** illustrates how structured noise can alter the correlation of noisy pattern estimates in a simple case of just two task conditions. Even if we assume the noise is spatially and temporally independent (i.e., **Σ*_ϵ_*** is a diagonal matrix, which may be a valid assumption if one “pre-whitens” the data before further analysis [25]), the bias structure still exists but reduces to **(X*^T^* X)^−1^*σ*^2^**, where ***σ*^2^** is the variance of the noise.

Since the covariance matrix of 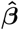 is biased, its correlation is also distorted from the true correlation structure. This is because correlation is merely a rescaling of rows and columns of a covariance matrix. Fig **1C** essentially illustrates this bias structure after being converted to correlation matrix (in this case, ***σ***=1 and ***β* = 0**) as this RSA structure, by virture of being derived for white noise, can only result from structure in the design matrix **X**. In reality, both spatial and temporal correlations exist in fMRI noise, which complicates the structure of the bias. But the fact that bias in Fig **1C** arises even when applying RSA to white noise which itself has no spatial-temporal correlation helps to emphasize the first contributor to the bias: the timing structure of the task, which is exhibited in the correlations between the regressors in the design matrix. Whenever the interval between events of two task conditions is shorter than the length of the HRF (which typically outlasts 12 s), correlation is introduced between their corresponding columns in the design matrix. The degree of correlation depends on the overlapping of the HRFs. If one task condition often closely precedes another, which is the case here as a consequence of the Markovian property of the task, their corresponding columns in the design matrix are more strongly correlated. As a result of these correlations, **X*^T^* X** is not a diagonal matrix, and neither is its inverse **(X*^T^* X)^−1^**.

In general, unless the order of task conditions is very well counter-balanced and randomized across participants, the noise **(X*^T^* X)^−1^X*^T^ ϵ*** in 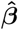 is not i.i.d between task conditions. The bias term **B = (X*^T^* X)^−1^X*^T^* Σ*_ϵ_*X(X*^T^* X)^−1^** then deviates from a diagonal matrix and causes unequal distortion of the off-diagonal elements in the resulting correlation matrix of 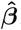. These unequal distortions alter the order of ranking of the values of the off-diagonal elements. Therefore, rank correlation between the similarity matrix from traditional RSA and similarity matrix of any candidate computational model is necessarily influenced by the bias. Conclusion based on such comparison between two similarity matrices or based on comparing a pair of off-diagonal elements within a neural similarity matrix becomes problematic, as long as the bias causes unequal distortion. Furthermore, if the design matrices also depend on participants’ performance such as errors and reaction time, the bias structure could depend on their performance as well. Comparison between neural representational structure and participants’ behavioral performance may also become problematic in such situations.

It is worth pointing out that the bias is not restricted to using correlation as metric of similarity. Because structured noise exists in 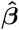, any distance metrics between rows of 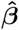 estimated within imaging runs of fMRI data are likely biased. We can take Euclidean distance as an example. For any two task conditions *i* and *j*, the expectation of the distance between 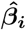 and 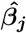 is 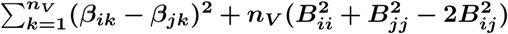, where **B** is the bias in the covariance structure. Therefore, the bias 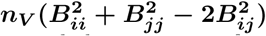 in Euclidean distance also depends on the task timing structure and the property of noise. (See Fig **1D**).

In our derivations above, point estimates of 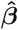 introduce structured noise due to the correlation structure in the design matrix. One might think that the bias can be avoided if a design matrix is not used, i.e., if RSA is not performed after GLM analysis, but directly on the raw fMRI patterns. Such an approach still suffers from bias, for two reasons that we detail below.

First, RSA on the raw activity patterns suffers from the second contributor to the bias in RSA that comes from the temporal properties of fMRI noise. To understand this, consider that estimating activity pattern by averaging the raw patterns, for instance 6 sec after each event of a task condition (that is, at the approximate peak of the event-driven HRF) is equivalent to performing an alternative GLM analysis with a design matrix **X_6_** that has delta functions 6 sec after each event. Although the columns of this design matrix **X_6_** are orthogonal and 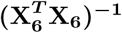 becomes diagonal, the bias term is still not a diagonal matrix. Because of the autocorrelation structure **Σ*_ϵ_*** in the noise, the bias term 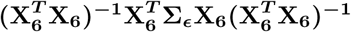 essentially becomes a sampling of the temporal covariance structure of noise at the distances of the inter-event intervals. In this way, timing structure of the task and autocorrelation of noise together still cause bias in the RSA result.

To illustrate this, we applied RSA to the raw patterns of an independent set of resting state fMRI data from the Human Connectome Project [26], pretending that the participants experienced events according to the 16-state task in Fig **1A**. As shown in Fig **1E**, even in the absence of any task-related signal spurious similarity structure emerges when RSA is applied to the raw patterns of resting state data. To quantify the extent of spurious structure in Fig **1E**, we computed the null distribution of the average estimated similarity structure by randomly permuting the task condition labels on each simulated participant’s estimated similarity structure 10000 times and averaging them. We then compared the absolute values of the off-diagonal elements in Fig **1E** against those in the null distribution. The Bonferroni corrected threshold for incorrectly rejecting at least one true hypothesis that an off-diagonal element in the average similarity matrix is from the null distribution is p=0.0004 for ***α***=0.05. In our resting-state fake RSA matrix, 39 out of 120 off-diagonal elements significantly deviated from the null distribution based on this threshold.

Second, averaging raw data 6 sec after events of interest over-estimates the similarity between neural patterns of adjacent events, an effect independent of the fMRI noise property. This is because the true HRF in the brain has a protracted time course regardless of how one analyzes the data. Thus the estimated patterns (we denote by 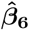) in this approach are themselves biased due to the mismatch between the implicit HRF that this averaging assumes and the real HRF. The expectation of 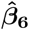 becomes 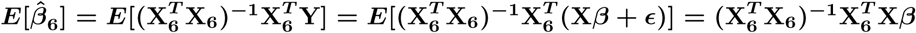 instead of ***β***. Intuitively, X temporarily smears the BOLD patterns of neural responses close in time but 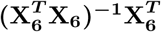 only averages the smeared BOLD patterns without disentangling the smearing. 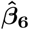 thus mixes the BOLD activity patterns elicited by all neural events within a time window of approximately 12 sec (the duration of HRF) around the event of interest, causing over-estimation of the similarity between neural patterns of adjacent events. If the order of task conditions is not fully counter-balanced, this method would therefore still introduce into the estimated similarity matrix a bias caused by the structure of the task.

Similar effect can also be introduced if 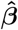 is estimated with regularized least square regression [27]. Regression with regularization of the amplitude of 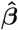 trades off bias in the estimates for variance (noise). On the surface, reducing noise in the pattern estimates may reduce the bias introduces into the similarity matrix. However, the bias in 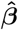 itself alters the similarity matrix again. For example, in ridge regression, an additional penalization term ***λβ^T^ β*** is imposed for ***β*** for each voxel. This turns estimates 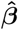 to 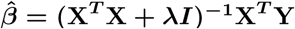. The component contributed to 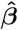 by the true signal ***Xβ*** becomes **(X*^T^* X + *λI*)^−1^X*^T^ β***. As ***λ*** increases, this component increasingly attributes neural activity triggered by other task events near the time of an event of interest to the event’s activity. Therefore, this method too would overestimate pattern similarity between adjacent events.

In all the derivations above, we have assumed for simplicity of illustration that the noise in all voxels has the same temporal covariance structure. In reality, the autocorrelation can vary over a large range across voxels (Fig. **1G**). So the structured noise in each voxel would follow a different distribution. Furthermore, the spatial correlation in noise means the noise in 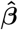 is also correlated across voxels, which makes the bias even more complicated. At minimum, noise correlation between voxels violates the requirement of Pearson correlation that pairs of observations should be independent.

### Bayesian RSA significantly reduces bias in the estimated similarity

As shown above, the covariance structure of the noise in the point estimates of neural activity patterns 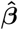 leads to bias in the subsequent similarity measures. The bias can distort off-diagonal elements of the resulting similarity matrix unequally if the order of task conditions is not fully counterbalanced. In order to reduce this bias, we propose a new strategy that aims to infer directly the covariance structure ***U*** that underlies the similarity of neural patterns, using raw fMRI data. Our method avoids estimating 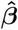 altogether, and instead marginalizes over the unknown activity patterns ***β*** without discarding uncertainty about them. The marginalization avoids the structured noise introduced by the point estimates, which was the central cause of the bias. Given that the bias comes not only from the experimental design but also from the spatial and temporal correlation in noise, we explicitly model these properties in the data. We name this approach Bayesian RSA (BRSA) as it is an empirical Bayesian method [28] for estimating ***U*** as a parameter of the prior distribution of ***β*** directly from data.

### Direct estimation of similarity matrix while marginalizing unknown neural patterns

BRSA assumes a hierarchical generative model of fMRI data. In this generative model, the covariance structure ***U*** serves as a hyper-parameter that governs the distribution of ***β***, which in turn generates the observed fMRI signal **Y**. Each voxel ***i*** has its own noise parameters, including auto-correlation coefficient ***ρ_i_***, variance 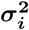 of innovation noise (the noise component unpredictable from the previous time step) and pseudo-SNR ***s_i_*** (we use the term ‘pseudo-SNR’ because the actual SNR depends on both the value of the shared covariance structure ***U*** and the voxel-specific scaling factor ***s_i_***). Given these, **(*σ_i_s_i_*)^2^*U*** is the covariance matrix of the distribution of the activity levels ***β_i_*** in voxel ***i***. The model allows different signal and noise parameters for each voxel to accommodate situations in which only a fraction of voxels in an ROI might have high response to tasks [27] and because the noise property can vary widely across voxels (e.g., Fig. **1G**). We denote the voxel-specific parameters (**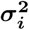*, ρ_i_*** and ***s_i_***) of all voxels together as ***θ***.

If the fMRI noise can be assumed to be independent across voxels [19], then for any single voxel ***i***, we can marginalize over the unknown latent variable ***β_i_*** to obtain an analytic form of the likelihood of observing the fMRI data ***Y_i_*** in that voxel ***p*(*Y_i_*|X*, U, θ_i_*)**. Multiplying the likelihoods for all voxels will result in the likelihood for the entire dataset: ***p*(Y|X*, U, θ*)**. Note that this computation marginalizes over ***β***, avoiding altogether the secondary analysis on the point estimates 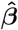 that is at the heart of traditional RSA. Through the marginalization, all the uncertainty about ***β*** is correctly incorporated into the likelihood. By searching for the optimal 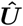 and other parameters 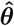 that maximize the data likelihood, we can therefore obtain a much less biased estimate of ***U*** for the case of spatially independent noise [19].

However, as illustrated by [14], intrinsic fluctuation shared across brain areas that is not driven by stimuli can dominate the fMRI time series and influence the RSA result. If one labels any fluctuation not captured by the design matrix as noise, then intrinsic fluctuation shared across voxels can manifest as spatial correlation in the noise, which violates our assumption above. To reduce the impact of intrinsic fluctuation on the similarity estimation, we therefore incorporate this activity explicitly into the BRSA method, with inspiration from the GLM denoising approach [29].

We start by assuming that the shared intrinsic fluctuation across voxels can be explained by a finite set of time courses, which we denote as ***X*_0_**, and the rest of the noise in each voxel is spatially independent. If ***X*_0_** were known, the modulation ***β*_0_** of the fMRI signal **Y** by ***X*_0_** can be marginalized together with the response amplitude ***β*** to the experimental design matrix ***X*** (note that we still infer ***U***, the covariance structure of ***β***, not of ***β*_0_**). Since ***X*_0_** is unknown, BRSA uses an iterative fitting procedure that alternates between a step of fitting the covariance structure ***U*** while marginalizing ***β*_0_** and ***β***, and a step of estimating the intrinsic fluctuation ***X*_0_** from the residual noise with principal component analysis (PCA). Details of this procedure are described in the Materials and Methods under *Model fitting procedure*.

Since our goal is to estimate ***U***, voxel-specific parameters ***θ*** can also be analytically or numerically marginalized so that we only need to fit ***U*** for the marginal likelihood ***p*(Y|X, X_0_*, U*)**. This reduces the number of free parameters in the model and further allows for the extension of estimating a shared representational structure across a group of participants, as shown later. Fig **2** shows a diagram of the generative model. More details regarding the generative model and the marginalization can be found in the Materials and Methods, under *Generative model of Bayesian RSA*.

**Figure 2.**
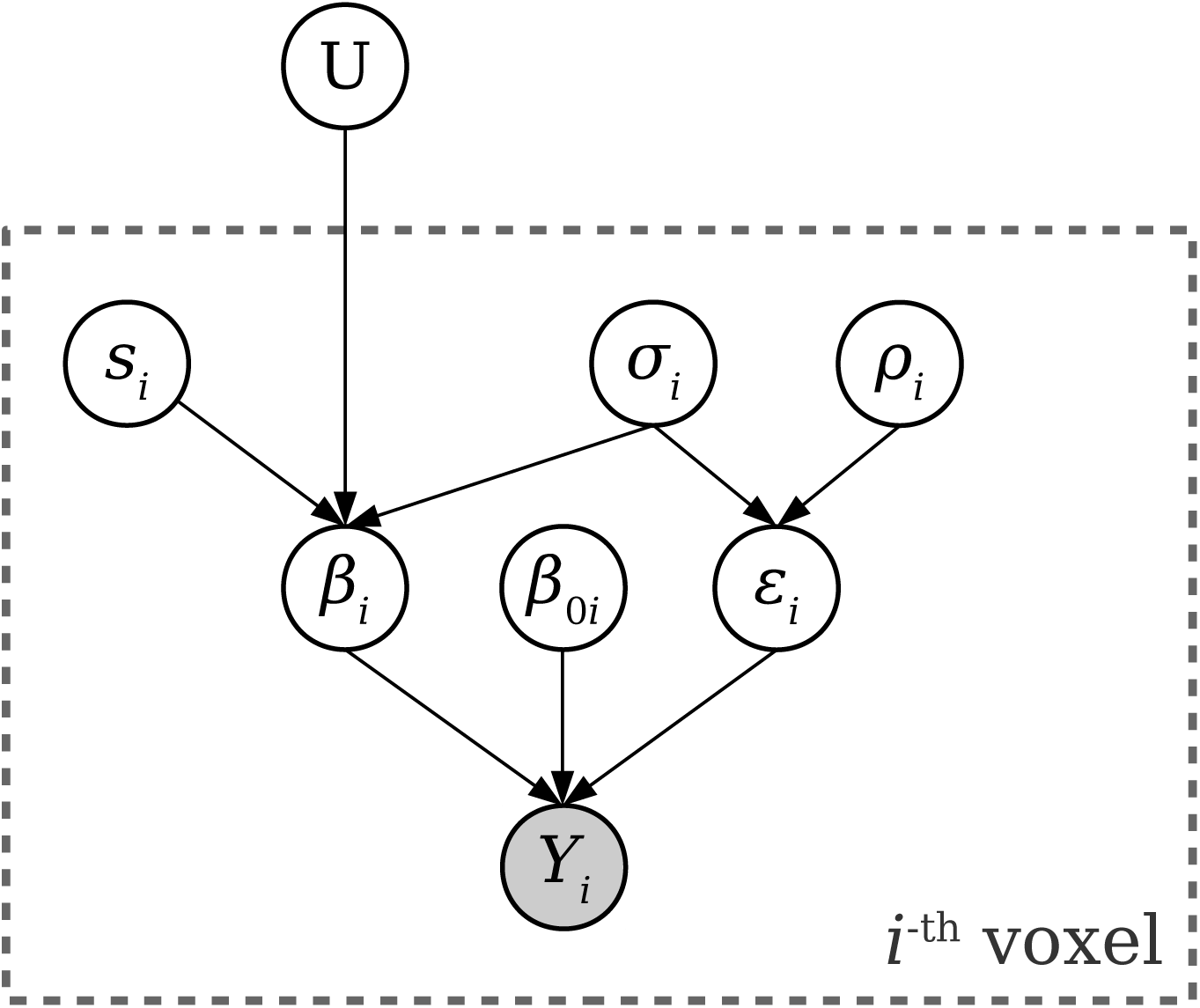
Generative model of Bayesian RSA. The covariance structure ***U*** shared across all voxels is treated as a hyper-parameter of the unknown response amplitude *β*. For voxel ***i***, the BOLD time series ***Y_i_*** are the only observable data. We assume ***Y_i_*** is generated by task-related activity amplitudes ***β_i_*** (the ***i***-th column of ***β***), intrinsic fluctuation amplitudes ***β*_0*i*_** and spatially independent noise ***ϵ_i_***: ***Y_i_* = X*β_i_* + X_0_*β*_0*i*_ + *ϵ_i_***, where **X** is the design matrix and **X_0_** is the set of time courses of intrinsic fluctuations. ***ϵ_i_*** is modeled as an AR(1) process with autocorrelation coefficient ***ρ_i_*** and noise standard deviation ***σ_i_***. ***β_i_*** depends on the voxel’s pseudo-SNR ***s_i_*** and noise level ***σ_i_*** in addtion to ***U***: ***β_i_* ∼ *N*(0, (*s_i_σ_i_*)^2^*U*)**. By marginalizing over ***β_i_***, ***β*_0*i*_**, ***σ_i_***, ***ρ_i_*** and ***s_i_*** for each voxel, we can obtain the likelihood function ***p*(*Y_i_*|X, X_0_*, U*)** and search for ***U*** which maximizes the total log likelihood 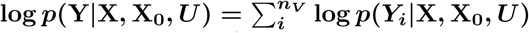 of the observed data **Y** for all ***n_V_*** voxels. The optimal 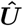 can be converted to a correlation matrix, representing the estimated similarity between patterns.

The covariance matrix ***U*** can be parameterized by its Cholesky factor ***L***, a lower-triangular matrix. To find the 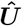 that best explains the data ***Y***, we first calculate the 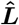 that best explains the data by optimizing the marginal log likelihood:
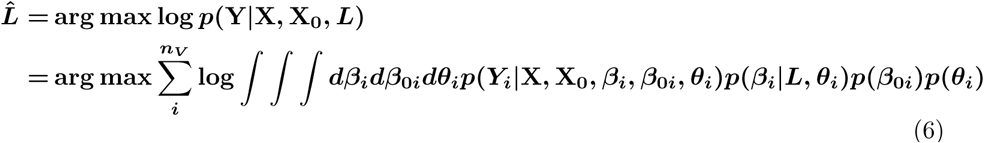

And then obtain the estimated covariance matrix
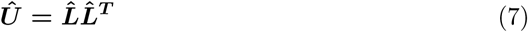

Once 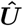 is estimated (after the iterative fitting procedure for ***L*** and ***X*_0_**), 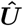 is converted to a correlation matrix to yield BRSA’s estimation of the similarity structure.

### BRSA recovers simulated similarity structure

To test the performance of BRSA in a case where the ground-truth covariance structure is known, we embedded structure into resting state fMRI data. Signals were simulated by first sampling response amplitudes according to a hypothetical covariance structure for the “16-state” task conditions (Fig **3A**), and then weighting the design matrix of the task in Fig **1A** by the simulated response amplitudes. The simulated signals were then added to resting state fMRI data. In this way, the “noise” in the test data reflected the spatial and temporal structure of realistic fMRI noise. To make the estimation task even more challenging, we simulated a situation in which within the ROI (Fig **3B**; we took the lateral occipital cortex as an ROI in this simulation, as an example) only a small set of voxels respond to the task conditions (Fig **3C**). This is to reflect the fact that SNR often varies across voxels and that an ROI is often pre-selected based on anatomical criteria or independent functional localizer, which do not guarantee that all the selected voxels will have task-related activity.

**Figure 3.**
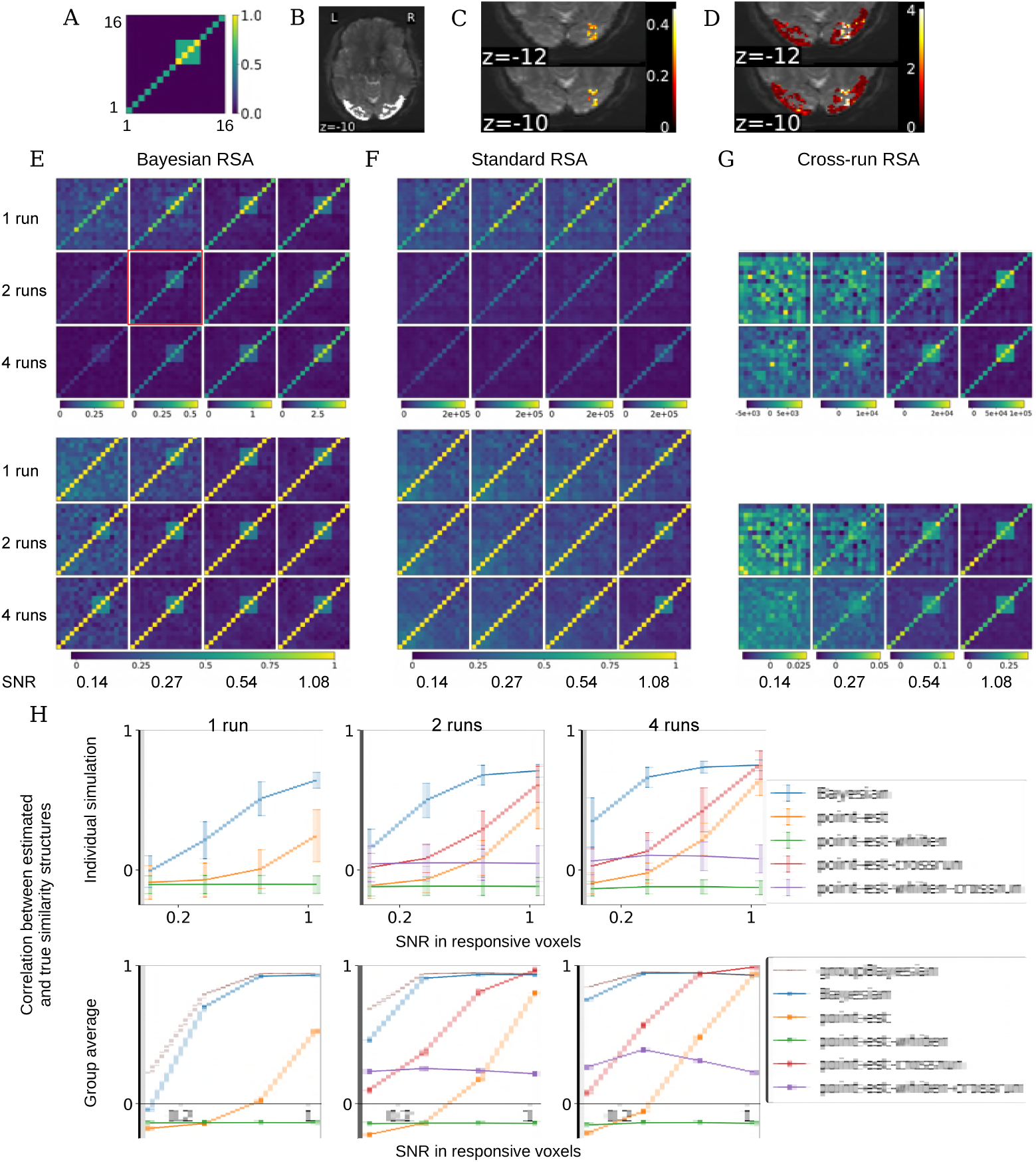
Performance of BRSA on simulated data. **(A)** The true covariance structure ***U*** from which the activity patterns were drawn. **(B)** We use lateral occipital cortex (bright region) as an example ROI and resting state fMRI data from the Human Connectome Project as noise. **(C)** We multiplied the design matrix of the task in Fig 1A with the simulated activity pattern and then added this “signal” to voxels that in a cubical region of the ROI. The colors show the actual SNR of the added signal for one example simulated brain, corresponding to the plot circumvented by a red square in E. **(D)** The pseudo-SNR map estimated by BRSA for the data with a true SNR map shown in C. The scale does not match the scale of true SNR, but the spatial pattern of SNR is recovered. **(E)** Average covariance matrix (top) and similarity matrix (bottom) estimated by BRSA in the cubic area in C, across different SNR levels (columns) and different numbers of runs (rows). **(F)** The corresponding result obtained by standard RSA based on activity patterns estimated within runs. **(G)** The corresponding result of RSA based on cross-correlating patterns estimated from separate runs. **(H)** Top: average correlation (mean **±** std) between the off-diagonal elements of the estimated and true similarity matrices, for each method, across SNR levels (x-axis) and amounts of data (separate plots). Bottom: The correlation between the average estimated similarity matrix of each method (for GBRSA, this is the single similarity matrix estimated) and the true similarity matrix. “point-est”: methods based on point estimates of activity patterns; “-crossrun”: similarity based on cross-correlation between runs; “-whiten”: patterns were spatially whitened (similarity matrix not shown because the true structure could barely be seen)

Fig **3E** shows the average covariance structure and similarity matrix estimated by BRSA. The corresponding results estimated based on 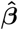 in standard RSA are shown in Fig **3F**. This comparison clearly demonstrates that at low SNR and with a small amount of data, BRSA can recover the simulated covariance structure of task-related signals, while standard RSA is overwhelmed by the bias structure (eq. 5). It has been suggested that cross-run RSA, that is, similarity calculated between patterns estimated from separate scanning runs, can also reduce bias [14, 15, 17]. As shown in Fig **3G**, indeed the true covariance and similarity structure can be recovered better by this approach as compared to within-run RSA (Fig **3F**). However, this approach leads to faster degradation of results as SNR decreases, as demonstrated by the lowest two SNR levels in the simulation. The peak height of task-triggered response is often in the range of 0.1-0.5% in cognitive studies [30] while the noise level is often a few percents, which means the SNRs expected in real studies are likely in the lower range in our simulation, except when studying primary sensory stimulation. Furthermore, the inner products or correlation between noises in patterns estimated from separate runs can be positive or negative by chance. When the noise is large enough, even the correlation between pattern estimates in different runs corresponding to the same task conditions may become negative (as observed in Fig **3G**). This makes it difficult to associate results of cross-run RSA with a notion of pattern “similarity” because one would not expect patterns for a task condition to be anti-correlated across runs. Fig **3H** summarizes the average correlation between the off-diagonal elements of the estimated similarity matrix and those of the simulated similarity matrix. At high SNR, cross-run RSA’s performance is similar to that of BRSA, and they both outperform within-run RSA. But BRSA performs the best at low SNR.

We also tested cross-run RSA with the estimated patterns spatially whitened using the procedure of [17]. Surprisingly, spatial whitening hurts similarity estimation. This might be because the spatial correlation structure of the simulated signal is different from that of the noise. Whitening based on the spatial correlation structure of noise would re-mix signals between different voxels to the extent of changing its similarity structure. Practically, it is difficult to estimate the spatial correlation of true signal patterns, because their estimates are always contaminated by noise.

### Added bonus: inferring pseudo-SNR map

Although the voxel-specific parameters ***θ*** are marginalized during fitting of the model, we can obtain their posterior distribution and estimate their posterior means. The estimated pseudo-SNR 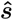 is of particular interest, as it informs us of where the estimated representational structure is more strongly supported in the ROI chosen by the researcher. As shown in Fig **3D**, the estimated pseudo-SNR map highly resembles the actual map of SNR in our simulated data in Fig **3C**, up to a scaling factor.

### Estimating shared representational similarity across participants

As mentioned above, BRSA can be extended to jointly fit the data of a group of participants, thus identifying the shared representational similarity structure that best explains the data of all participants. This is achieved by searching for a single ***U*** that maximizes the joint probability of observing all participants’ data (Group Bayesian RSA;GBRSA). The rationale or GBRSA is that it searches for the representational structure that best explains all the data. Using all the data to constrain the estimation of ***U*** reduces the variance of estimation for individual participants, an inspiration from hyper-alignment [20] and shared response model [21]. Fig **3H** shows that the similarity structure recovered by GBRSA has slightly higher correlation with the true similarity structure than the average similarity structure estimated by other methods, across most of the SNR levels and amounts of data. Cross-run RSA performs better only at the highest simulated SNR. However, low average SNR is common in many brain areas and this is where (G)BRSA offers more power for detecting the true but weak similarity structure.

### Controlling for over-fitting: model selection by cross-validation on left-out data

Although Fig **2** shows that BRSA reduces bias, it does not eliminate it completely. This may be due to over-fitting to noise. Because it is unlikely that the time course of intrinsic fluctuation ***X*_0_** and the design matrix ***X*** are perfectly orthogonal, part of the intrinsic fluctuation cannot be distinguished from task-related activity. Therefore, the structure of ***β*_0_**, the modulation of intrinsic fluctuation, could also influence the estimated 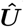 when SNR is low.

For instance, in Fig **3E**, at the lowest SNR and least amount of data (top left subplot), the true similarity structure is almost undetectable using BRSA. Is this due to large variance in the estimates, or is it because BRSA is still biased, but to a lesser degree than standard RSA? If the result is still biased, then averaging results across subjects will not remove the bias, and the deviation of the average estimated similarity structure from the true similarity structure should not approach 0. To test this, we simulated many more subjects by preserving the spatial patterns of intrinsic fluctuation and the auto-regressive properties of the voxel-specific noise in the data used in Fig **3**, and generating intrinsic fluctuations that maintain the amplitudes of power spectrum in the frequency domain. To expose the limit of the performance of BRSA, we focused on the lower range of SNR and simulated only one run of data per “subject”. Fig **4A** shows the quality of the average estimated similarity matrix with increasing number of simulated subjects. The average similarity matrices estimated by BRSA do not approach the true similarity matrix indefinitely as the number of subjects increase. Instead, their correlation saturates to a value smaller than 1. This indicates that the result of BRSA is still weakly biased, with the bias depending on the SNR. It is possible that as the SNR approaches 0, the estimated 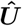 is gradually dominated by the impact of the part of ***X*_0_** not orthogonal to ***X***. We reason that this is partly because the algorithm [31] we used to estimate the number of components in ***X*_0_** is a relatively conservative method. In particular, in this simulation, the number of components of simulated intrinsic fluctuations were 20**±**4, while the number of components estimated from these simulated data by the algorithm were 13**±**3. However, empirically this algorithm [31] yields more stable and reasonable estimation than other methods we have tested [32]. It should be noted that BRSA still performs much better than standard RSA, for which the correlation between the estimated similarity matrix and the true similarity matrix never passed 0.1 in these simulations (not shown).

**Figure 4.**
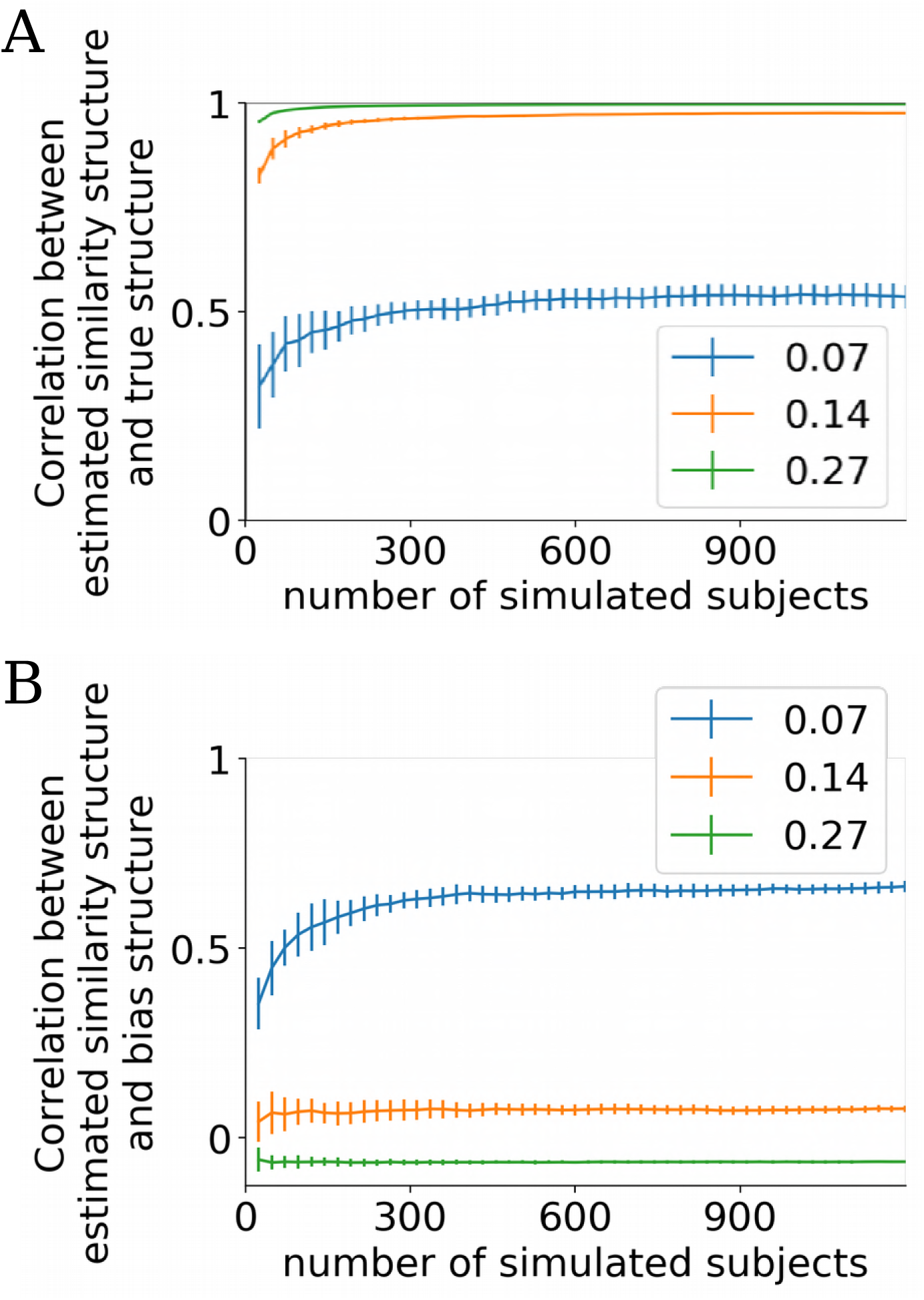
Limited performance of BRSA at very low SNR and small amount of data. **(A)** The average correlation between the off-diagonal elements of the estimated and true similarity matrices (mean **±** std) as the number of simulated subjects increases. Each simulated subject had one run of data. Legend shows average SNR in task-responsive voxels. Half of the voxels do not include any signal related to the design matrix. The correlation reaches asymptotic levels slightly below 1 with increasing numbers of participants except when the SNR is extremely low (0.07), indicating that the bias is not fully eliminated.**(B)** The average correlation between the estimated similarity matrix and the expected bias structure assuming white noise. The estimated similarity structure is most dominated by the bias structure at the lowest SNR simulated (0.07). The negative correlation at the highest SNR reflects the weak negative correlation between the true similarity structure and expected bias structure (−0.055)

The expected bias structure when spatial noise correlation exists is difficult to derive. We used **(X*^T^* X)^−1^** as a proxy to evaluate the residual bias in the estimated similarity using BRSA. As expected, when the SNR approached zero, the model over-fit to the noise and the bias structure increasingly dominated the estimated structure despite increasing the number of simulated participants (Fig **4B**). This observation calls for an evaluation procedure to detect over-fitting in applications to real data, when the ground truth of the similarity structure is unknown.

One approach to assess whether a BRSA model has over-fit the noise is cross-validation. In addition to estimating ***U***, the model can also estimate the posterior mean of all other parameters, including the neural patterns ***β*** of task-related activity, ***β*_0_** of intrinsic fluctuation, noise variances ***σ*^2^** and auto-correlation coefficients ***ρ***. For a left-out testing data set, the design matrix ***X_test_*** is known given the task design. Together with the parameters estimated from the training data as well as the estimated variance and auto-correlation properties of the intrinsic fluctuation in the training data, we can calculate the log predictive probability of observing the test data. The unknown intrinsic fluctuation in the test data can be marginalized by assuming their statistical property stays unchanged from training data to test data. The predictive probability can then be contrasted against the cross-validated predictive probability provided by a null model separately fitted to the training data. The null model would have all the same assumptions as the full BRSA model, except that it would not assume any task-related activity captured by ***X***. When BRSA over-fits the data, the estimated spatial pattern 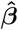 would not reflect the true response pattern to the task and is unlikely to be modulated by the time course in ***X_test_***. Thus the full model would predict signals that do not occur in the test data, and yield a lower predictive probability than the null model. The result of the full BRSA model on training data can therefore be accepted if the log predictive probability by the full model is higher than that of the null model significantly more often than chance.

Over-fitting might also arise when the assumed design matrix ***X*** does not correctly reflect task-related activity. When there is a sufficient amount of data but the design matrix does not reflect the true activity, the estimated covariance matrix 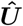 in BRSA would approach zero, as would the posterior estimates of 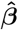. In this case as well, the full model would be indistinguishable from the null model.

We tested the effectiveness of relying on cross-validation to reject over-fitted results using the same simulation procedure as in Fig **3**, and repeated this simulation 36 times, each time with newly simulated signals and data from a new group of participants in HCP [33] as “noise”. Fig **5A** shows the rate of correct acceptance when both training and test data have signals. We counted each simulation in which the cross-validation score (log predictive probability) of the full BRSA model was significantly higher than the score of the null model (based on a one-sided student’s t-test at a threshold of ***α***=0.05) as one incidence of correct acceptance. When the SNR is high (above 0.14), warranting reliable estimation of the similarity structures as indicated in Fig **3H**, the cross-validation procedure selected the full model significantly more often than chance (all p<7e-7, binomial test). At the lowest SNR (0.14) and with only 1 run of training data, the full model was never selected (p<3e-11), consistent with a poor estimation of the similarity matrix in Fig **3E**. As the amount of training data increased to 2 runs (even without changing the SNR), the rate of accepting the full model increased, although with the lowest SNR it was still not significantly different from chance (p=0.6), while the estimated similarity matrix was also noisy but started to be visually detectable in Fig **3E**. This indicates that the cross-validation procedure is relatively conservative. Fig **5B** shows the difference between the cross-validation scores of the full and null models as the amount of training data doubled in one group of simulated subjects, as an illustration of typical results. Dots to the left of the dashed line represent subjects for whom the full model explained the data better than the null model to a greater extent when two runs were used, as compared to one run of data only. The means and standard deviations of the t-statistics across simulated groups for all simulation configurations are displayed in Fig **5C**. The differences in cross-validation scores between full and null models are displayed in Fig **5D**.

**Figure 5.**
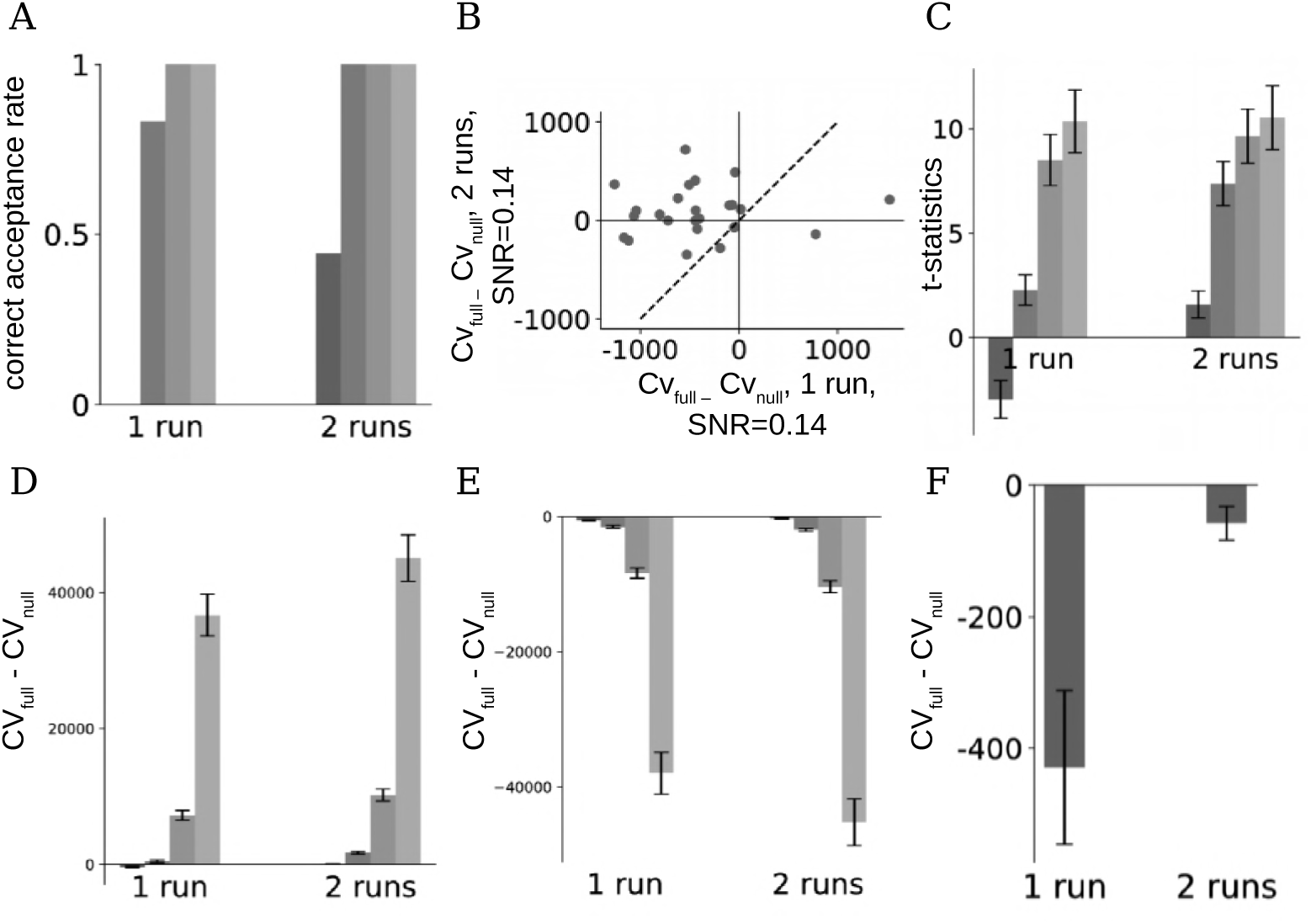
Cross-validation reduces the chance of false positive results. The full BRSA model and a null model that assumes no task-related activity were fit to 1 or 2 runs of simulated data of a group of subjects with different SNRs, as in Fig 3. Student’s t-test was performed on the difference between the cross-validation scores of the full model and null model on a left-out run of data to determine whether the full model should be accepted. This procedure was repeated 36 times on different groups of simulated data. **(A)** Signals were added to both training and test data. The frequencies with which the full models were accepted based on the t-test (correct acceptance) are displayed. Darker bars correspond to low SNRs and lighter bars correspond to higher SNRs in Fig 3. **(B)** The difference between cross-validation scores of the full and null models for 1 or 2 runs of training data, at the lowest SNR (0.14). Dashed line: ***x* = *y***. Points to the left of the dashed line show more evidence for the full model when two runs of data were used, as compared to one run. The chance of accepting the full model increases when there are more data to fit the BRSA model. **(C)** Mean **±** std of the t-statistics of the difference between cross-validation scores of the full and null models across simulated groups, for the corresponding amounts of data and SNR in A. **(D)** Mean **±** std of the difference between the cross-validation scores of the full models and the null models across simulated groups in A. **(E)** Mean **±** std of the difference between the cross-validation scores when only the training data but not test data have signals. **(F)** Mean **±** std of the difference between the cross-validation scores when neither the training data nor the test data have signals. In all cases in E and F the statistical test correctly rejected the full model.

The cross-validation procedure also helps avoid false acceptance when activity patterns are not consistently reproducible across runs. To illustrate this, we simulated the case when signals are only added to the training data but not to the test data. Now, the full model was always rejected across the simulated SNR and amounts of data (not shown). Finally, when neither training data nor testing data included signal, the cross-validation procedure also correctly rejected the full model in all cases. Fig **5E** and **5F** illustrate the difference between cross-validation scores of full and null models for the two simulations, respectively.

### Extension: decoding task-related activity from new data

BRSA has a relatively rich model for the data: it attempts to model both the task-related signal and intrinsic fluctuation, and to capture voxel-specific SNR and noise properties. In addition to cross-validation, this also enables decoding of signals related to task conditions from new data. Similarly to the procedure of calculating cross-validated log likelihood, but without pre-assuming a design matrix for the test data, we can calculate the posterior mean of 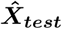 and 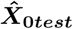 in the testing data. Fig **6A** shows the decoded design matrix 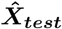 for one task condition (condition 6 in Fig **1B** and **6B**) and one participant, using one run of training data with the second-highest SNR. Although our method decodes some spurious high responses when there is no event of this task condition, overall the result captures many of the true responses in the design matrix. The average correlation between the decoded design matrix and the true design matrix is displayed in Fig **6B**. High values on the diagonal elements indicate that overall, the decoder based on BRSA can recover the task-related signals well. The structure of the off-diagonal elements appears highly similar to those of the correlation structure between corresponding columns in the original design matrix(r=0.82, p<1e-30). This means that the signals corresponding to task conditions which often occur closely in time in training data are more likely be confused when they are decoded from testing data. Indeed, at the time of mistakenly decoded high response around the 90th TR in Fig **6A**, there is a true event of the first task condition ((Fo)Fy) in the design matrix. The decoder confused the response to the first condition as response to the sixth condition. The events of these two conditions did in fact often co-occur in the training data, therefore their overlap in the design matrix makes it difficult to distinguish which event triggered the response in the training data, and reduces the accuracy of posterior estimates of their activity patterns, causing further confusion at the stage of decoding. We suspect that such confusion is not limited to decoding based on BRSA, but should be a general limitation of multi-variate pattern analysis of fMRI data: due to the slow smooth BOLD response, the more often the events of two task conditions occur closely in time in the training data, the more difficult it becomes for the classifier to discern their patterns.

**Figure 6.**
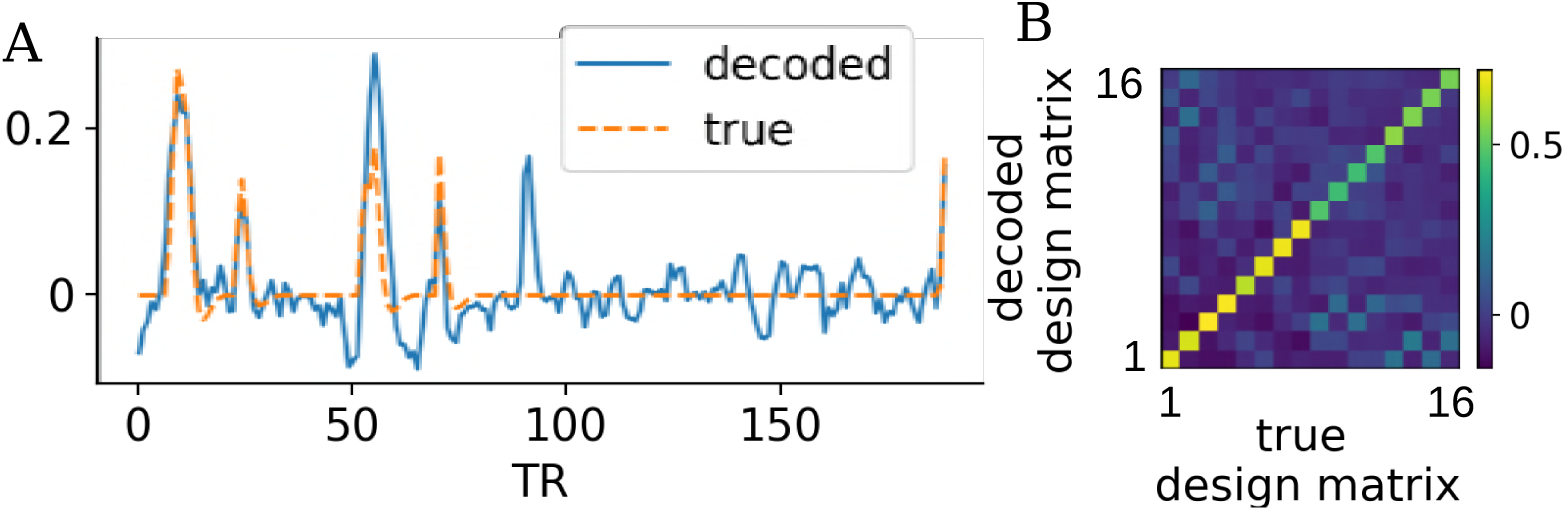
Decoding capabilities of the BRSA method. **(A)** Decoded task-related activity of the sixth condition from one simulated subject in one run of test data, and the true design matrix of that condition in the test data. The simulated data with the second highest SNR in Fig 3 were used. BRSA model was fitted to one run of training data. **(B)** Average correlation between the decoded signals for each task condition (rows) and the time courses for each condition in the design matrix used to simulate the test data (columns).

## Discussion

In this paper, we demonstrated that bias can arise in the result of representational similarity analysis, a popular method in many recent fMRI studies. By analytically deriving the source of the bias with simplifying assumptions, we showed that it is determined by both the timing structure of the experiment design and the correlation structure of the noise in the data. Traditional RSA is based on point estimates of neural activation patterns which unavoidably include high amounts of noise. The task design and noise property induce covariance structure in the noise of the pattern estimates. This structure in turn biases the covariance structure of these point estimates, and a bias persists in the similarity matrix. Such bias is especially severe when the SNR is low and when the order of the task conditions cannot be fully counterbalanced.

To reduce this bias, we proposed a Bayesian framework that interprets the representational structure as reflecting the shared covariance structure of activity levels across voxels. Our BRSA method estimates this covariance structure directly from data, bypassing the structured noise in the point estimates of activity levels, and explicitly modeling the spatial and temporal structure of the noise. This is different from many other methods that attempt to correct the bias after it has been introduced.

In addition to inferring the representational similarity structure, our method also infers activation patterns (as an alternative to the traditional GLM), SNR for different voxels, and even the “design matrix” for data recorded without knowledge of the underlying conditions. The inferred activation patterns are regularized not only by the SNR, but also by the learned similarity structure. The inference of an unknown “design matrix” allows one to uncover uninstructed task conditions (e.g., in free thought) using the full Bayesian machinery and all available data.

In a realistic simulation using real fMRI data as background noise, we showed that BRSA generally outperforms standard RSA and cross-run RSA, especially when SNR is low and when the amount of data is limited, making out method a good candidate in scenarios of low SNR and difficult-to-balance tasks. Because temporal and spatial correlation also exist in the noise of data from other neural recording modalities, the method can also be applied to other types of data when the bias in standard RSA is of concern. To detect overfitting to noise, the difference between the cross-validated score of the full model of BRSA and a null model can serve as the basis for model selection. We further extend the model to allow for estimating the shared representational structure across a group of participants.

The bias demonstrated in this paper does not necessarily question the validity of all previous results generated by RSA. However, it does call for more caution when applying RSA to higher-level brain areas for which SNR in fMRI is typically low, and when the order between events of different task conditions cannot be fully counterbalanced. This is especially the case with decision making tasks that involve learning or structured sequential decisions, in which events cannot be randomly shuffled. Even when the order of task conditions can be randomized, it may not be perfectly counter-balanced. Thus, a small deviation of the bias structure from a diagonal matrix may still exists. If the same random sequence is used for all participants, the tiny bias can persist in the results of all participant and become a confound. Therefore, it is also important to use different task sequences across participants.

Prior to the proposal of our method, similarity measures calculated between patterns estimated from separate scanning runs (cross-run RSA) was proposed to overcome the bias [15, 17, 34]. The inner product between noise pattern estimates from separate runs is theoretically unbiased. However, at low SNR, cross-run RSA suffers from large noise, sometimes generating results where noisy pattern estimates of the same condition from different runs appear anti-correlated. In addition, even though the cross-run covariance matrix is not biased, the magnitude of cross-run correlation is under-estimated because the computation requires division by the standard deviation of the estimated patterns, which is in turn inflated by the high amount of noise carried in the estimated patterns. In our simulation, cross-run RSA appears to slightly outperform BRSA at very high SNR but the results of both methods are already very close to the true similarity structure in this case. On the other hand, at very low SNR, cross-run RSA fails to reveal the true similarity structure, while BRSA does. However, cross-run RSA may be a more conservative approach given that the cross-run covariance matrix (and Mahalonobis distance [17] is unbiased. It is difficult to predict whether BRSA or cross-run RSA are more suitable for any specific study and brain area of interest. Nonetheless, based on our results, both approaches should always be favored over traditional within-run RSA based on pattern estimates.

It is surprising that spatial whitening, that is often recommended [17], in fact hurts the result of standard RSA and cross-run RSA in our simulation. This may be because, in our simulation, the spatial correlation of noise is not the same as the spatial correlation in the simulated signal. While whitening reduces the correlation between noise in the estimated 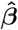 of different voxels, it may cause undesired remixing of true signals between voxels. As discussed before, in practice, it is difficult to know whether the intrinsic fluctuation and task-evoked signals share the same spatial correlation structure, because we do not know the ground truth of signals in real data. The cost and benefit of spatial whitening on standard and cross-validated RSA therefore awaits more studies. Instead of performing spatial whitening, BRSA estimates a few time series ***X*_0_** that best explain the correlation of noise between voxels and marginalizes their modulations in each voxel. Without remixing signals across voxels, it still captures spatial noise correlation.

In our study, we did not directly compare BRSA to cross-validated Mahalanobis distance [17] because they are foundamentally different measures: BRSA aims to estimate the correlation between patterns, which is close to the cosine angle between two patterns vectors [35, 36]; in contrast, Mahalanobis distance aims to measure the distance between patterns. Nonetheless, given the theoretical soundness of the cross-validated Mahalanobis distance, it could also be a good alternative to BRSA when there are multiple runs in a task.

Our BRSA method is closely related to the PCM [16, 37]. A major difference is that PCM models the point estimates 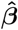 after GLM analysis while BRSA models fMRI data **Y** directly. The original PCM [16] in fact considered the contribution of the noise in pattern estimates to the similarity matrix, but assumed that the noise in 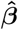 is i.i.d across task conditions. This means that the bias in the covariance matrix was assumed to be restricted to a diagonal matrix. We showed here that when the order of task conditions cannot be fully counter-balanced, such as in the example in Fig **1**, this assumption is violated and the bias cannot be accounted for by methods such as PCM.

If one knew the covariance structure of the noise **Σ*_ϵ_***, then the diagonal component of the noise covariance structure assumed in PCM [16] could be replaced by the bias term **(X*^T^* X)^−1^X*^T^* Σ*_ϵ_*X(X*^T^* X)^−1^** to adapt PCM to estimate the covariance structure 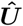 that best explains 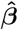 [38] if spatial noise correlation is not considered. However, as shown in Fig **1G**, different voxels can have a wide range of different autocorrelation coefficients. Assuming a single **Σ*_ϵ_*** for all voxels may be over-simplified. In addition, PCM assumes all voxels within one ROI have equal SNR. However, typically only a small fraction of voxels exhibits high SNR [27]. Therefore, it is useful to model the noise property and SNR of each voxel individually.

In addition to these differences, BRSA explicitly models spatial noise correlation. It also comes with the ability to select between a full model and null model based on cross-validated log likelihood, and the method can be applied to fMRI decoding. PCM can additionally evaluate the likelihood of a few fixed candidate representational structures given by different computational models. It can also estimate the additive contributions of several candidate pattern covariance structures to the observed covariance structure. These options are not yet available in the current implementation of BRSA. Combining the strength of PCM and BRSA is an interesting future direction.

Many aspects of flexibility may be incorporated to BRSA. For example, the success of the analysis hinges on the assumption that the HRF used in the design matrix correctly reflects the true hemodynamics in the ROI, but it has been found that HRF in fact vary across people and across brain regions [39, 40]. Jointly fitting the shape of the HRF and the representational structure may improve the estimation. In addition, it is possible that even if the landscape of activity patterns for a task condition stays the same, the global amplitude of the response pattern may vary across trials due to repetition supression [41–43] and attention [44, 45]. Such modulation may not be predictable by response time or stimulus duration. Allowing global amplitude modulation of patterns associated with a task condition to vary across trials might capture such variability and increase the power of the method.

Our simulations revealed that BRSA is not entirely unbiased, that is, results cannot be improved indefinitely by adding more subjects. We hypothesize that the residual bias is due to the underestimation of the number of components necessary to capture the spatial correlation introduced by intrinsic fluctuation. Development of a proper but less conservative algorithm for estimating the number of components suitable for BRSA may improve its performance.

Comparing the cross-validation score of the full model and a null model is one approach to detect overfitting. One interesting finding is that when the design matrix does not explain the real brain response (Fig **5C** where signal was not added to either training or test data), and when there is a sufficient amount of training data, the full model becomes indistinguishable from the null model. Even though such cross-validation does not select the null model significantly more often than the full model, not finding the opposite is sufficient to warn the researcher not to trust the resulting similarity matrix as reflecting the true structure. When this happens, it is advisable to focus on taking measures to improve the design of study. Ultimately, task designs that are not fully counterbalanced and low SNR in fMRI data are two critical factors that cause bias in traditional RSA and impact the power of detecting similarity structure. Carefully designing tasks that fully balance the task conditions, randomizing the sequence of a task across participants, and increasing the number of measurements, are our recommended approaches in the first place. In the analysis phase of the project, one can then use BRSA.

## Materials and methods

### Generative model of Bayesian RSA

Our generative model of fMRI data follows the general assumption of GLM. In addition, we model spatial noise correlation by a few time series ***X*_0_** shared across all voxels. The contribution of ***X*_0_** to the ***i^th^*** voxel is ***β*_0*i*_**. Thus, for voxel ***i***, we assume that
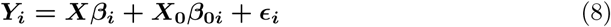

***Y_i_*** is the time series of voxel ***i***. ***X*** is the design matrix shared by all voxels. ***β_i_*** is the response amplitudes of the voxel ***i*** to the task conditions. ***ϵ_i_*** is the residual noise in voxel ***i*** which cannot be explained by either ***X*** or ***X*_0_**. We assume that ***ϵ*** is spatially independent across voxels, and all the correlation in noise between voxels are captured by the shared intrinsic fluctuation ***X*_0_**.

We use an AR(1) process to model ***ϵ_i_***: for the ***i^th^*** voxel, we denote the noise at time ***t >* 0** as ***ϵ_t,i_***, and assume
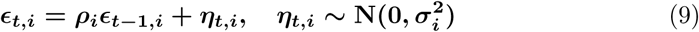

where 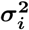 is the variance of the “innovation” noise at each time point and *ρ_i_* is the autoregressive coefficient for the *i^th^* voxel.

We assume that the covariance of the multivariate Gaussian distribution from which the activity amplitudes ***β_i_*** are generated has a scaling factor that depends on its pseudo-SNR ***s_i_***:
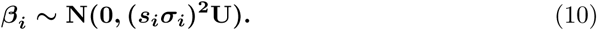

This is to reflect the fact that not all voxels in an ROI respond to tasks.

We further use Cholesky decomposition to parametrize the covariance structure ***U : U*** = ***LL^T^***, where ***L*** is a lower triangular matrix. Thus, ***β_i_*** can be written as ***β_i_* = *s_i_σ_i_Lα_i_***, where ***α_i_*** ∼ ***N*(0*, I*)**. This change of parameter allows for estimating **U** of lower rank (if the researcher has sufficient reason to make such a guess) by setting ***L*** as lower-triangular matrix with a few rightmost-columns truncated. With an improper uniform prior for ***β*_0*i*_**, and temporarily assuming ***X*_0_** is given, we have the unmarginalized likelihood for each voxel ***i***:
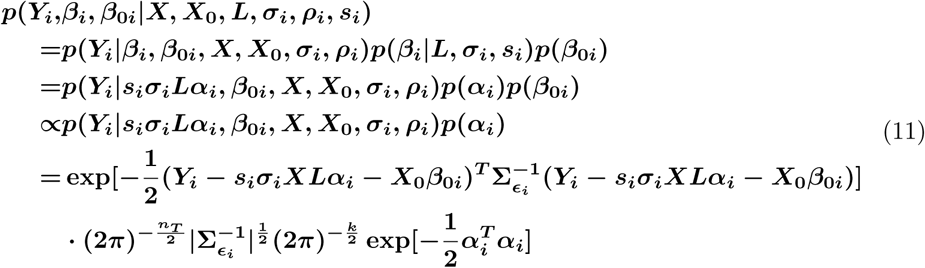

where ***k* ≤ *n_C_*** is the rank of ***L***.

In contrast to the full model, our null model assumes
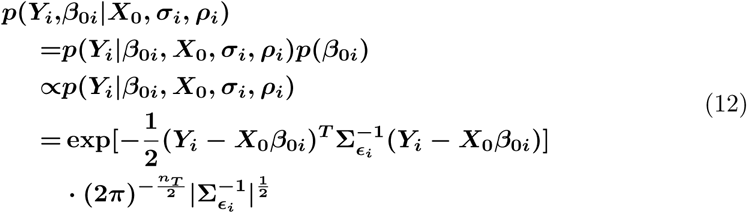

For data within one run, 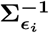, the inverse matrix of the covariance of ***ϵ_i_***, is a banded symmetric matrix which can be written as 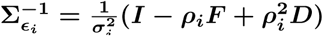, where *F* is 1 only at the superdiagonal and subdiagonal elements and 0 everywhere else, and ***D*** is 1 on all diagonal elements except for the first and last one, and 0 elsewhere. For abbreviation, we can denote 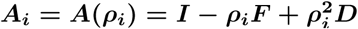 which is a function of ***ρ_i_***. 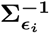 can be factorized as 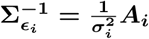.When *Y_i_* includes concatenated time series across several runs, 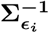 is a block diagonal matrix with each block diagonal elements corresponding to one run, constructed in the same way.

To derive the log likelihood of ***L*** for data of all voxels in the ROI, we need to marginalizing all other unknown parameters. Below, we marginalize them step by step.

By marginalizing ***β*_0*i*_**, we have
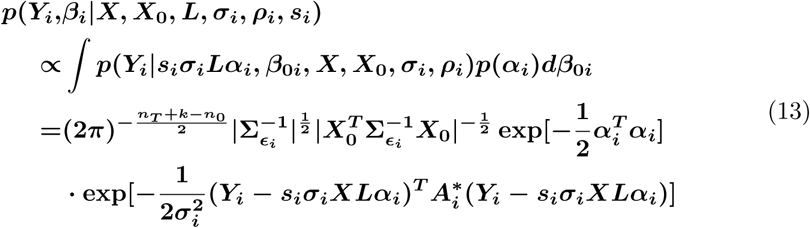

***n*_0_** is the number of components in ***X*_0_**. In the equation above, we denoted 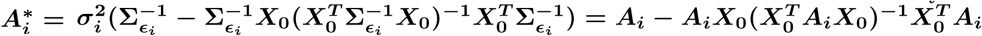.

By further marginalizing ***α_i_*** which is equivalent to marginalizing ***β_i_***, we get
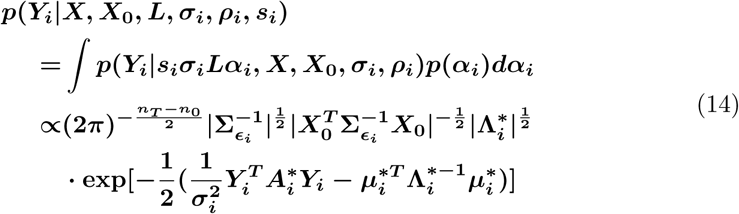

where 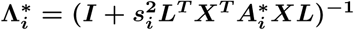 and 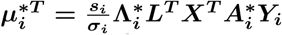 are the variance and mean of the posterior distribution of ***α_i_***, respectively.

All the steps of marginalization above utilize the property of multivariate Gaussian distribution. Next we marginalize the noise variance 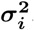. We assume an improper uniform distribution of 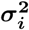 in 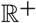. It is also possible to assume a conjugate prior for 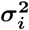. Given that data of at least hundreds of time points are obtained in each run to provide enough constraint to 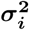, our choice does not appear to cause problem. To isolate 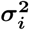, using the property of Cholesky decomposition of 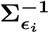, the above equation can be written as
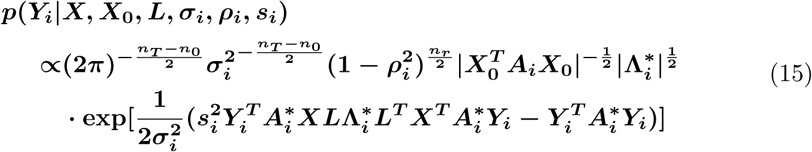

This form is proportional to an inverse-Gamma distribution of 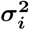. ***n_r_*** is the number of runs in the data. Therefore, we can analytically marginalize 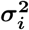 and obtain
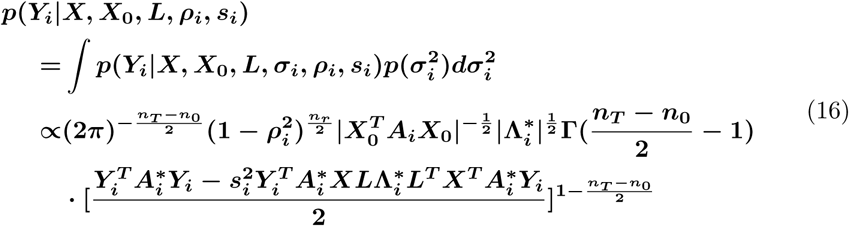

We did not find ways to further analytically marginalize ***s_i_*** or ***ρ_i_***. But we can numerically marginalize them by weighted sum of 16 at ***n_l_* × *n_m_*** discrete grids **{*ρ_il_, s_im_*}** (**0 *< l < n_l_***, **0 *< m < n_m_***) with each grid representing one area of the parameter space of **(*ρ, s*)**.
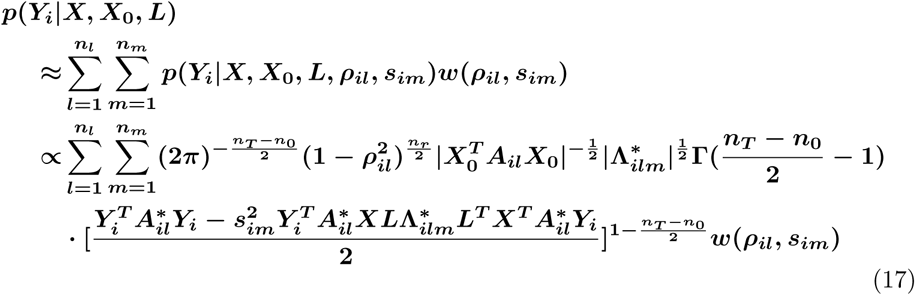

The weights ***w*(*ρ_il_, s_im_*)** are the prior probabilities of the two parameters in the area represented by **{*ρ_il_, s_im_*}**. We assume uniform prior of ***ρ*** in (−1,1). All the simulations in this paper used a negative exponential distribution as prior for ***s***. The grids ***s_im_*** are each chosen at the centers of mass of the prior distribution in the bins they represent in **(0, +∞)**. All bins equally divide the area under the curve of the prior distribution for ***s***. Alternative forms of priors such as uniform in (0, 1) and truncated log normal distribution are also implemented in the tool.

Because we made the assumption that ***ϵ_i_*** is independent across voxels. The log likelihood for all data is the sum of the log likelihood for each voxel.
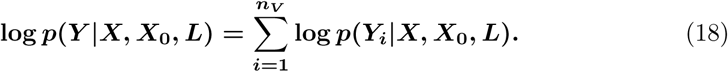

For the null model, the likelihood for each voxel after marginalizing ***β*_0*i*_** and 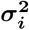 can be similarly derived,
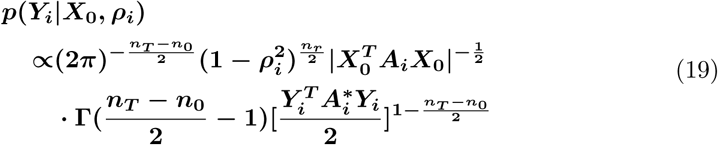

and the total log likelihood can be calculated similarly by numerically marginalizing ***ρ_i_*** and summing the log likelihood for all voxels.

### Model fitting procedure

To fit the model, we need the derivative of the total log likelihood with respect to ***L***. It can be derived that conditional on any grid of parameter pairs **{*ρ_il_, s_im_*}**, the derivative of the log likelihood for voxel ***i*** against each lower-triangular element of ***L*** is the corresponding lower-triangular element of the matrix
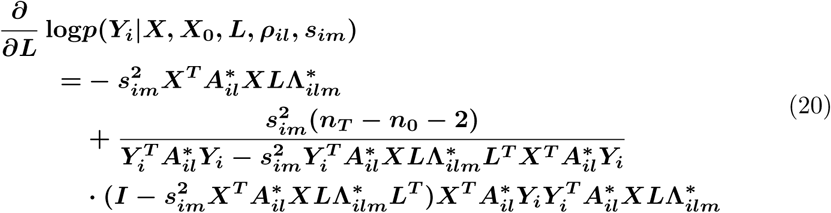

where 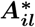 and 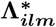 are 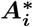 and 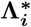 evaluated at {*ρ_il_*,*s_im_*}. The derivative of the total log likelihood against L after marginalizing over all grids **{*ρ_il_, s_im_*}** of all voxels is
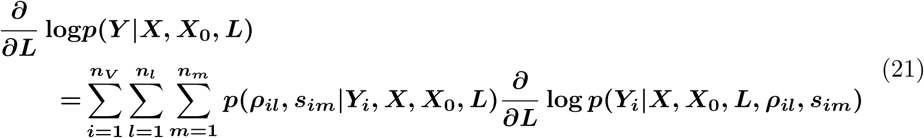

***p*(*ρ_il_, s_im_*|*Y_i_, X, X*_0_*, L*)** is the posterior probability of **{*ρ_il_, s_im_*}** conditional on a given ***L***. It can be obtained by normalizing ***p*(*Y_i_*|*X, X*_0_*, L, ρ_il_, s_im_*)*w*(*ρ_il_, s_im_*)** after calculating 16.

With the derivative 21, the total log likelihood 18 can be maximized using gradient-based method such as Broyden–Fletcher–Goldfarb–Shanno (BFGS) algorithm to search for the optimal ***L*** [46–49].

However, the derivations above have made the assumption that ***X*_0_** is given, while it is not. The requirement for ***X*_0_** should be to appropriately capture the correlation of noise across voxels without overfitting. Therefore, at the starting of the model fitting, regular regression of ***Y*** against ***X*** and any nuisance regressors such as head motion and constant baseline is performed. Then the algorithm by Gavish and Donoho [31] is used to select the optimal number of components ***n*_0_** to choose ***X*_0_** from the eigenvectors of the residual of regression. Because regular regression does not shrink the magnitudes of ***β***, their magnitudes can only be over-estimated. ***n*_0_** thus has no risk of being over-estimated. This ***n*_0_** is then fixed throughout the model fitting. Next, the first ***n*_0_** principal components of the residual of regression are set as 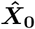 to allow for calculating the marginal log likelihood in 21 and gradient ascent with BFGS. A sufficient steps of iterations are performed to optimize ***L***. Then 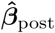, the posterior expectations of ***β***, are calculated with the current 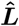 and with ***s***, ***ρ***, ***σ*** being marginalized. 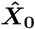 is subsequently recalculated using PCA from the residuals after subtracting 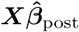 from ***Y***. The alternation between optimizing ***L*** and re-estimating 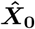 is repeated until convergence.

Once we obtain 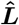, the estimate of ***L***, the estimate of the covariance structure is 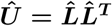. Converting it into a correlation matrix yields the similarity matrix by BRSA. Even though 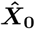 is estimated from data based on posterior estimation of ***β*** repeatedly during fitting, ***L*** is still optimized for the log likelihood with all other unknown variables marginalized. Thus the estimated 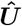 is an empirical prior of ***β*** estimated from data. This is the reason we consider our model as an empirical Bayesian method.

Many subcomponents of the expressions in these equations do not depend on ***L*** and thus can be pre-computed before optimizing for ***L***. The fixed grids of **(*ρ, s*)** further make several subcomponents shared across voxels when evaluating 16. These all reduce the amount of computation needed.

The fitting of the null model is similar to that of the full model except that there is no ***L*** to be optimized.

### Model selection and decoding

Once a model has been fitted to some data from a participant or a group of participants, we can estimate the posterior mean of ***ρ***, ***s***, ***σ*^2^**, ***β*** and ***β*_0_**, conditional on the empirical prior 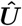 (essentially 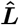), data ***Y***, design matrix ***X*** and estimated intrinsic fluctuations 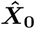. Below, we derive their formula and the procedure in which they are used for calculating cross-validated log likelihood of new data and decoding task-related signal 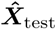 and 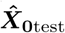 from new data in the context of fMRI decoding.

The posterior mean of these variables are
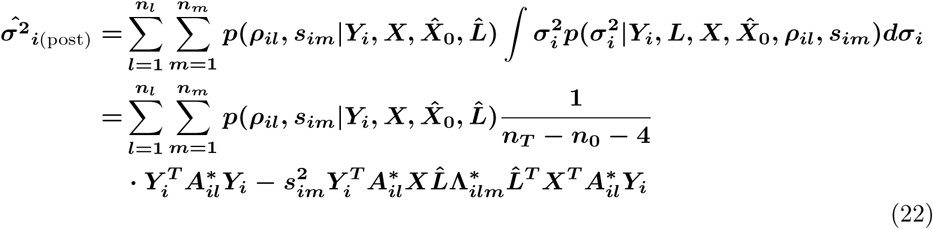

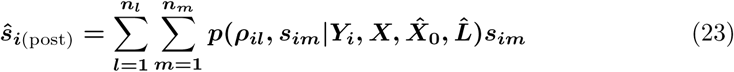

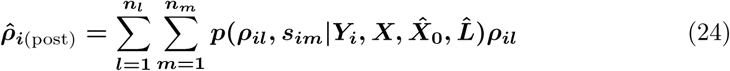

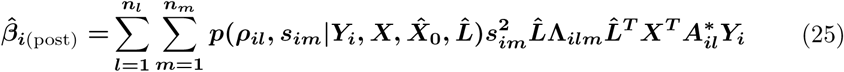

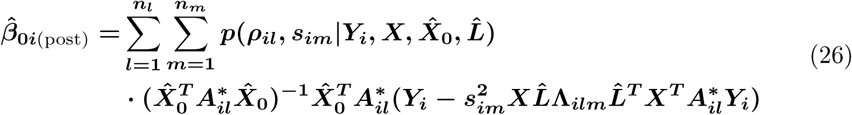

For null model, 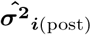, 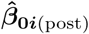 and 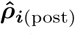 are of similar forms except that all terms including ***s_im_*** are removed and that 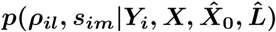 is replaced by 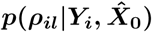.

To calculate cross-validated log likelihood, we assume the posterior estimates above and the statistical properties of ***X*_0_** stay unchanged in the testing data. We use zero-mean AR(1) process to describe the statistical properties of ***X*_0_**. The AR(1) parameters estimated from 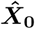 serve as the parameters of the empirical prior for ***X*_0_** in the testing data. When ***X*_0_** at each time point ***t*** is treated as a random vector 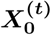, the AR(1) parameters of each component can be jointly written as the diagonal matrix 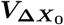 for the variance of the innovation noise, and diagonal matrix 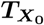 for the auto-regressive coefficients, both of size ***n*_0_ × *n*_0_**.

For model selection purpose, design matrix ***X***_test_ for the testing data should be generated in the same manner as they are for the training data by the researcher. For full BRSA model, 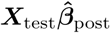 is the predicted task-related signal in ***Y***_test_.

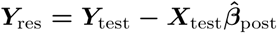 is the residual variation which cannot be explained by the design matrix and the posterior activity pattern 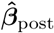. Null model does not predict any task-related activity, so all ***Y***_test_ constitutes residual variation ***Y***_res_. In either the full model or the null model, the posterior estimate 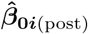 expresses their prediction about how voxels should be co-modulated by a fluctuation, while the fluctuation time course ***X*_0_**_test_ is only predictable in terms of its variance and temporal autocorrelation expressed by 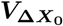 and 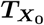. 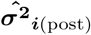 and 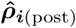 express the models’ predictions about the variance and temporal dependency of the fluctuation in each voxel in addition to the co-fluctuation. With these parameters estimated from training data, both the full and null models can marginalize the unknown ***X*_0_**_test_ and yield their corresponding predictive log likelihoods for the testing data ***Y***_test_. These log likelihoods are the basis for selecting between the full and null models.

To calculate the log likelihood, we notice that the predictive model of ***Y***_res_ in testing data by both models are dynamical system models in which ***X*_0_**_test_ is the latent state and ***Y***_res_ is the observed data. They are slightly different from the standard dynamical system model [50] in that not only the latent states, but also the noise, have temporal dependency [51]:
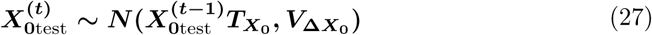

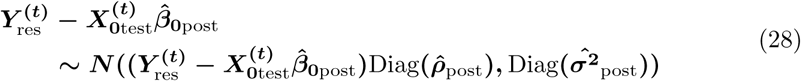

Where 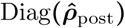 and 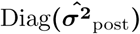 are diagonal matrices with vectors 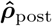 and 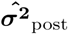 being their diagonal elements, respectively.

Because a modified forward-backward algorithm from the standard approach [50] is needed to calculate the preditive log likelihood 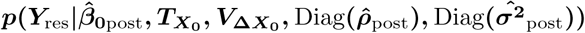 and the posterior distribution of ***X*_0_**_test_, we describe the procedure below.

Define
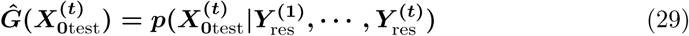

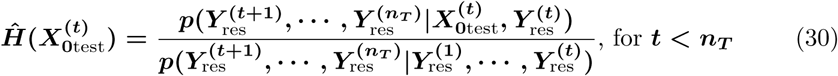

and 
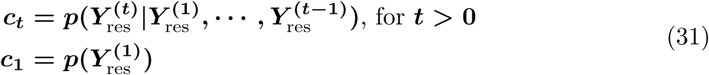

Therefore, the cross-validated log likelihood is
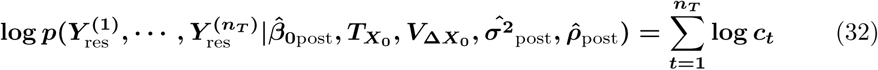

It can be derived that the posterior distribution of 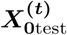 is
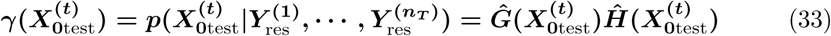

Below, we denote the mean and covariance of 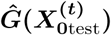 as 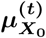 and 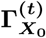, and the mean and covariance of 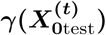 as 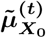 and 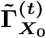.

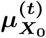, 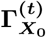 and *c_t_* can be calculated by the forward step. 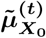 and 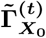 can be calculated by the backward step. To perform model selection, only forward step is necessary.

To perform the forward step, we first note that for *t* = 1
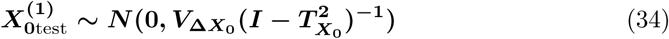

and 
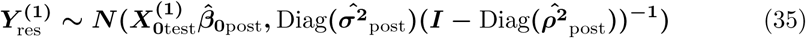

Denote 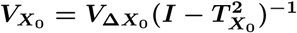, we have
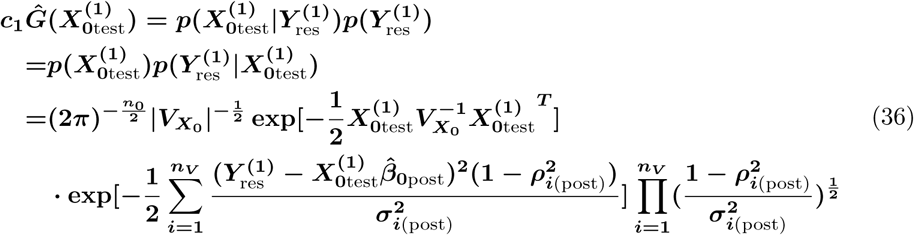

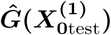 is a multivariate normal distribution of **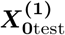,** we can find its covariance and mean from 36:
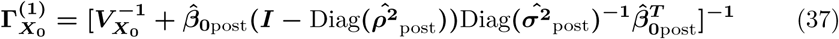

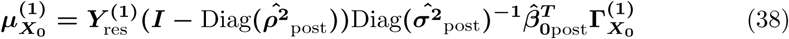

Because 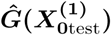 is a normalized probability distribution, the components in 36 after factoring out the multivariate normal distribution 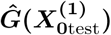 is ***c*_1_**:
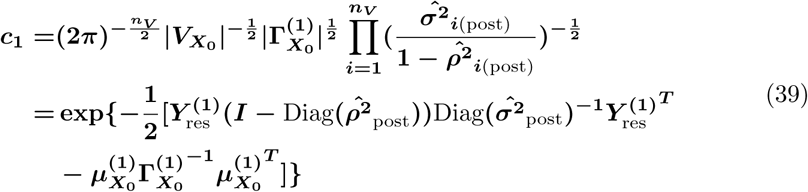

For any ***t >* 1**, the following relation holds:
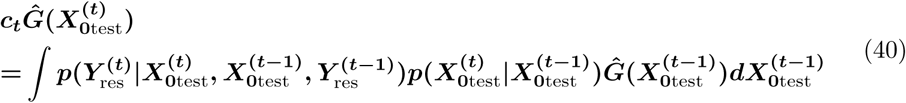

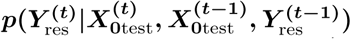 is defined by 28. 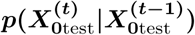 is defined by 27. Mean and covariance of 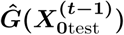 are calculated by the previous step for ***t* − 1**. Therefore, by marginalizing 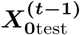, we obtain
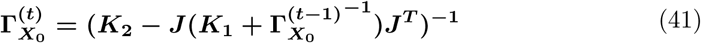

and 
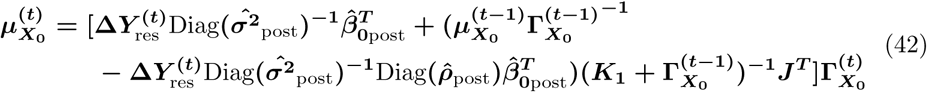

where **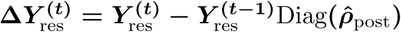.**

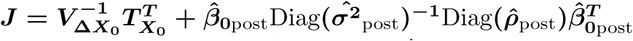, 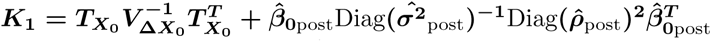 and 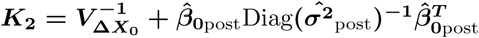. Note **J**, **K_1_**, **K_2_** are all constants.

Similarly to 39, after factoring out 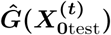, we obtain
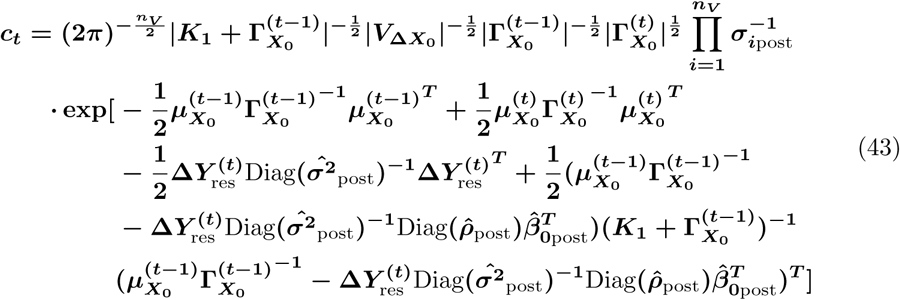

By calculating 41, 42 and 43 recursively with ***t*** incremented by 1 until ***n_T_***, the predictive log likelihood 32 of both the full and null models can be calculated to serve as the basis of model selection.

To calculate the mean and variance of the posterior distribution 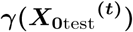 of ***X*_0_**_test_, backward step is needed. We denote its mean as 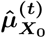, and covariance as 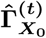.

For any ***t < n_T_***, it can be derived that
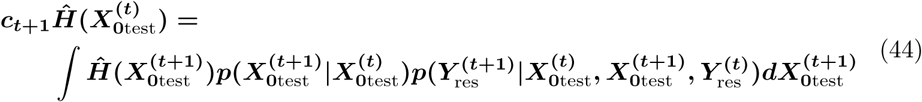

By plugging in 33, we get
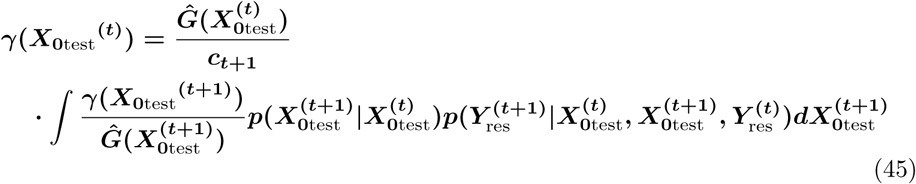

After the marginalization in 45 and observing the terms related to 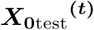, we get the recursive relations
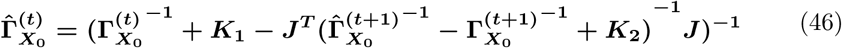

and
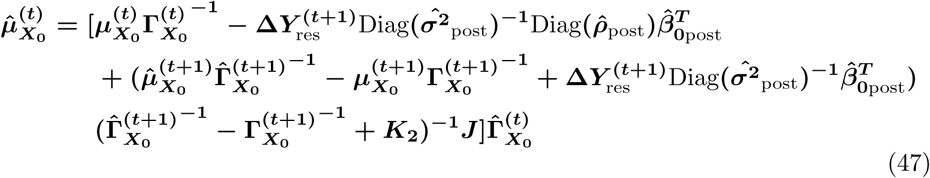

Note that **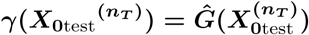,** therefore 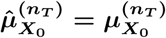 and 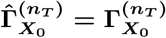. By recursively calculating 46 and 47 with ***t*** decremented by 1 from ***n_T_* − 1** until 1, the posterior distribution of 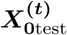 given all the testing data can be calculated.

For decoding purpose, we need to obtain not only the posterior mean of intrinsic fluctuations 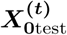, but also the task-related activity 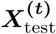. Therefore, we do not subtract a predicted signal 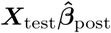 based on a hypothetical design matrix from testing data ***Y***_test_. We perform the forward-backward algorithm on ***Y***_test_ directly. By replacing 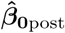 in the equations from 27 to 47 with 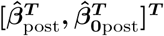 and other related terms accordingly, the posterior mean of both 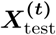 and 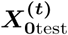 can be decoded just as 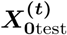 is decoded in 47.

### Data processing and analysis

Data used in Fig **1B** are from the experiments of Schuck et al. [22], following the same preprocessing procedure as the original study. The fMRI data were acquired at TR=2.4s. Data of 24 participants were used. Their design matrices were used for all the following analyses and simulations. Data in **Fig 1E,G** and Fig **3** were preprocessed data obtained from Human Connectome Project (HCP) [33]. The first 24 participants who have completed all 3T protocols and whose data were acquired in quarter 8 of the HCP acquiring period without image quality issues were selected for analysis in Fig 3. Data from 864 participants without image quality issues in HCP were used in the analysis in Fig 5. Each participants in the HCP data have 2 runs of resting state data with posterior-anterior phase encoding direction and 2 runs with anterior-posterior phase encoding direction. Time series were resampled at the same TR as the design matrix before further analysis.

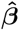 point estimates in Fig **1** were obtained with AFNI’s *3ddeconvolve* [52]. The design matrices were set up by convolving the stereotypical double-Gamma HRF in SPM [53] with event time courses composed with impulses lasting for the duration of the participants’ reaction time. AR(1) coefficients in Fig **1G** were estimated after upsampling the fMRI time series in the HCP data to the TR in Schuck et al. [22] and linear detrending. Upsampling is to reflect the lower temporal resolution more typically employed in task-related fMRI studies.

In the experiments of Fig **3**, lateral occipital cortex was chosen as the ROI, which included **4804 ± 29** (mean **±** standard deviation) voxels. Task related signals were only added to voxels within a bounding box of which the coordinates satisfy **25 < *x* < 35**, **−95 < *y* < −5** and **−15 < *z* < 5**. **189.0 ± 0.2** voxels fell within this bounding box. ***β*** were simulated according to the covariance matrix in Fig **3A** and scaled by one values in 1, 2, 4, 8. To evaluate the performance of the recovered correlation structure by different methods, the correlation between the off-diagonal elements of the recovered similarity matrix from data of each simulated participant was correlated with those elements of the ideal similarity matrix to yield the top panel of Fig **3H**. The top panel reflects the correlation of individual results. The bottom panel reflects the correlation of average results over simulated participants.

In order to make fair comparison with BRSA which considers temporal auto-correlation in noise, all the point estimates of 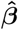 in Fig **3** were performed with restricted maximum likelihood estimation. AR(1) parameters of each voxel were estimated after initial regular regression. The AR(1) parameters were used to re-compute the temporal noise covariance matrices for each voxel and 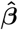 were calculated again assuming these noise covariance matrices. When spatial whitening of 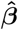 were performed, it followed the procedure of Diedrichsen et al. [17]. Point estimates of the spatial covariance of noise were first calculated from residuals of regression. These are not full rank matrices due to large numbers of voxels. The off-diagonal elements were further shrunk by weighting the point estimate of the spatial noise covariance structure by 0.4 and a diagonal covariance matrix with the same diagonal elements as the point estimate covariance matrix by 0.6.

To simulate the fMRI noise in Fig **4**, we first estimated the number of principal components to describe the spatial noise correlation in the 24 resting state fMRI data from HCP databse using the algoritm of Gavish and Donoho [31]. The spatial patterns of these principal components were kept fixed as the modulation magnitude ***β*_0_** by the intrinsic fluctuation. AR(1) parameters for each voxel’s spatially indepndent noise were estimated from the residuals after subtrating these principal components. For each simulated subject, time courses of intrinsic flucutations were newly simulated by scrambling the phase of the Fourier transformation of the ***X*_0_** estimated from the real data, thus preserving the amplitudes of their frequency spectrum. AR(1) noise were then added to each voxel with the same parameters as estimated from the real data. To speed up the simulation, only 200 random voxels from the ROI in Fig **3B** were kept for each participant in these simulations. Among them, 100 random voxels were added with simulated task-related signals. Thus, each simulated participant has different spatial patterns of ***β*_0_** due to the random selection of voxels. 500 simulated datasets were generated based on the real data of each participant, for each of the three SNR levels. In total 36000 subjects were simulated. The simulated pool of subjects were sub-divided into bins with a fixed number of simulated subjects ranging from 24 to 1200. The mean and standard deviation of the correlation between the true similarity matrix and the average similarity matrix based on the subjects in each bin were calculated, and plotted in Fig **4A**.

All SNRs in Fig **3** and Fig **4** were calculated post hoc, using the standard deviation of the added signals in the bounding box region devided by the standard deviation of the noise in each voxel, and averaged across voxels and simulated subjects for each level of simulation.

## Acknowledgments

This work was funded by the Intel Corporation and the John Templeton Foundation (15541). The opinions expressed are those of the authors and do not necessarily reflect the views of the John Templeton Foundation or of Intel Corporation. JWP was supported by grants from the McKnight Foundation, Simons Collaboration on the Global Brain (SCGB AWD1004351) and the NSF CAREER Award (IIS-1150186). Resting state fMRI data used in several experiments were obtained from the MGH-USC Human Connectome Project (HCP) database.

## Footnotes

1 Under the *brainiak.reprsimil.brsa* module. Our previous version of Bayesian RSA method [19] with newly added modeling of spatial noise correlation is in the *BRSA* class of the module. The new version described in this paper is implemented in the *GBRSA* class and can be applied to either a single participant or a group of participants.

